# Developing a Soft Micropatterned Substrate to Enhance Maturation of Human Induced Pluripotent Stem Cell-Derived Cardiomyocytes (hiPSC-CMs)

**DOI:** 10.1101/2024.07.12.599409

**Authors:** Yasaman Maaref, Shayan Jannati, Mohsen Akbari, Mu Chiao, Glen F Tibbits

## Abstract

Human induced pluripotent stem cell-derived cardiomyocytes (hiPSC-CMs) offer numerous advantages as a biological model, yet their inherent immaturity compared to adult cardiomyocytes poses significant limitations. This study addresses hiPSC-CM immaturity by introducing a novel physiologically relevant micropatterned substrate for long-term culture and maturation. A novel microfabrication technique combining laser etching and casting creates a micropatterned polydimethylsiloxane (PDMS) substrate with varying stiffness, from 2 to 50 kPa, mimicking healthy and fibrotic cardiac tissue, respectively. Platinum electrodes integrated into the cell culture chamber enabled pacing of cells at various frequencies. Subsequently, cells were transferred to the incubator for time-course analysis, ensuring contamination-free conditions. Cell contractility, cytosolic Ca^2+^ transient, sarcomere orientation, distribution, and nucleus aspect ratio are analyzed in a 2D hiPSC-CM monolayer up to 90 days post-replating in relation to substrate micropattern dimensions. Culturing hiPSC-CMs for three weeks on a micropatterned PDMS substrate (2.5-5 µm deep, 20 µm center-to-center spacing of grooves, 2-5 kPa stiffness) emerges as optimal for cardiomyocyte alignment, nucleus aspect ratio, contractility, and cytosolic Ca^2+^ transient. The study provides significant insights into substrate stiffness effects on hiPSC-CM contractility and Ca^2+^ transient at immature and mature states. Maximum contractility and fastest Ca^2+^ transient kinetics occur in mature hiPSC-CMs cultured for two to four weeks, with the optimum at three weeks, on a soft micropatterned PDMS substrate. This new substrate offers a promising platform for disease modeling and therapeutic interventions.

## 1. INTRODUCTION

Cardiovascular diseases are the leading cause of morbidity and mortality globally which account for 17.9 million deaths each year [1]. The heart is composed of various cell types, with cardiomyocytes emerging as the most dominant in terms of volume. In adult myocardial tissue, cardiomyocytes align into parallel cardiac muscle fibers characterized by highly oriented intercellular contractile myofibrils, which contain organized structures known as sarcomeres. Adult cardiomyocytes exhibit a rod-shaped morphology, characterized by highly aligned and organized sarcomeres with resting (diastolic) sarcomere lengths of about 2.2 µm [2]. They can be mono, bi-or poly-nucleated, and T-tubules are formed within their structure. This highly oriented structure is crucial for the proper excitation-contraction coupling of cardiomyocytes [3]. To gain a profound insight into cardiac disease pathogenicity, it is essential to establish a well-defined biological model. While adult cardiomyocytes obtained from explanted human hearts are suitable candidates for studying heart diseases, challenges arise due to limited sample availability, intricate pharmacological histories, limited viability in culture, no suitable controls, and preparation obstacles [4]. To address these limitations, hiPSC-CMs can be used as an unlimited cell source for cardiac disease modelling which provides novel insights into disease pathogenesis and sets the stage for testing novel therapeutic interventions [5][6]. However, in comparison to adult cardiomyocytes, immature hiPSC-CMs typically exhibit a more circular morphology, disorganized sarcomeres, mononucleation, and diffuse intercellular junctions [6][7]. Therefore, generating a physiologically relevant substrate for culturing hiPSC-CMs have been utilized to mitigate some of these limitations [5].

An ideal substrate for engineering functional 2D monolayer of cardiomyocytes needs to convey a blend of relevant mechanical and biological properties to facilitate cardiomyocyte (CM) attachment and growth, promote cellular alignment and elongation, and provide the flexibility for sustained cycles of expansion and contraction. Hence, the substrate stiffness should align with the stiffness of healthy cardiac tissue ranging from 1-6 kPa in the embryonic heart to approximately 10-15 kPa in the healthy adult heart [8]. Moreover, the substrate pattern should facilitate cell elongation and organization similar to that of adult cardiomyocytes. Thus, different microfabrication-based techniques, such as soft lithography, microcontact printing, and cast molding have been employed to create in vitro models for investigating the physiological and pathological characteristics of cardiomyocytes [9], [10].

Polydimethylsiloxane (PDMS) has been extensively used as a cell culture substrate, due to its high stability, excellent biocompatibility, tunable stiffness, gas permeability, transparency, and low fabrication cost, providing numerous advantages for cell growth [11]–[13]. PDMS can be readily fabricated and functionalized with extracellular matrix proteins such as fibronectin and laminin [14]. Additionally, PDMS does not exhibit poroelasticity, a characteristic observed in polyacrylamide (PA) gels, and it enables the creation of microscale patterns with high resolution [13]. Despite the numerous advantages of PDMS, their hydrophobic nature hinders the attachment of beating cardiomyocytes to the substrate, presenting a challenge in studying cell function in this physiologically relevant environment [15][16]. To reduce the surface hydrophobicity of PDMS substrates, different methods such as plasma treatment and ECM protein coating have been used [17]–[21]. However, the application of plasma-based treatment is restricted due to the hydrophobicity recovery, crack formation, and protein dissociation from the PDMS surface [22][23]. Previous studies have coated PDMS substrate with poly-dopamine (PD) to enhance the attachment and function of different cell types including mesenchymal stem cell, mouse embryonic stem cell, and hiPSCs [24]–[27].

Prior art creating soft culturing substrates for CMs includes the use of microcontact printing of extracellular matrix (ECM) proteins such as laminin and fibronectin onto PDMS and PA hydrogel [28]– [32]. The technique has been used to recapitulate the anisotropic alignment of CMs on substrates that mimic physiological stiffness. Microcontact printing is commonly employed for culturing individual cells or a line of cells, rather than for creating 2D monolayer cell formations [7], [29], [33]. Micromolding is an alternative technique that creates grooves and ridges serving as physical barriers and topographical cues to guide CM elongation and orientation [19]. The Khademhosseini group introduced an elastic micropatterned mechanical support through the photo crosslinking of methacrylate tropoelastin (MeTro) [17]. Their findings indicated that the dimension of the channels and the duration of culture significantly influenced the shape and orientation of actin filaments in CMs. In another study, a photodegradable Gelatin methacryloyl (PD GelMA) hydrogel was employed to make micropatterned cell culture substrates, which led to better alignment of cell nuclei along the channel direction and more regular beating rhythms [34]. However, the exploration of the impact of patterned substrates on CM nuclei alignment was restricted to days 1, 3, and 7, without extending the observation into the long term, spanning several weeks. This limitation may influence substrate topography, including alterations in the width and depth of the channels due to the swelling nature of PD GelMA hydrogel. This study also reported lower crosslink density at channel edges, causing local swelling, and changes in channel and groove dimensions.

Lind et al. [20] utilized a 3D printing technique to produce grooved microstructures using soft PDMS (SE1700, Dow-Corning) with a curing agent ratio of 1:25. Another technique used for generating grooves, ridges, microchannels and microfluidics is laser ablation of polymers and gelatins [21], [35], [36]. Nawroth et al. [37] introduced a laser-etching approach to create microscale surface groove and pillar structures, with feature dimensions in the order of 10–30 µm. They established a protocol for activating gelatin hydrogels using riboflavin-5’ phosphate, a non-toxic Ultraviolet (UV) photosensitizer. This process enabled the creation of micropatterns on the surface using a UV laser engraver. This method proved to be easier and faster in fabricating micropatterned substrates compared to traditional techniques involving photolithography and micro-molding. Chaofan et al. employed high-precision 3D printing to produce a mold for transferring the patterns onto gelatin-based hydrogels. Moreover, numerous studies have demonstrated the effectiveness of soft patterned substrates for single-cell analysis [17], [19]–[21], [35]–[37]. However, the reported cell culturing substrates typically exhibit stiffness ranging from 5 to 100 kPa, suitable for short cell culture durations of up to one week. Previous studies have only utilized a soft micropatterned substrate for single cell culture, not for 2D monolayer culture [38]. Moreover, the fabrication process of micropatterned substrates in these studies involved a clean room facility, making it a complex procedure [10]. Consequently, there exists a significant gap in the literature regarding the development of a straightforward protocol for employing soft substrates for extended periods, such as several weeks, and utilizing common biocompatible materials like PDMS. Notably, there has been no investigation focused on developing a soft micropatterned substrate with a stiffness of 2-5 kPa for the long-term culture (> 60 days) of spontaneously beating hiPSC-CMs in a 2D monolayer.

In this study, a novel microfabrication technique was employed to create a micropatterned substrate with physiologic and pathologic stiffness with a range of 2-5 kPa (soft) to 50 kPa (stiff) to recapitulate healthy and fibrotic human heart tissue, respectively. This fabrication process is straightforward and cost-effective compared to previous methods that require a clean room facility. To promote the maturation of hiPSC-CMs and recapitulate the physiological environment of cells, two key factors were considered during substrate fabrication: substrate stiffness and substrate micropatterning. To support the long-term culturing of 2D monolayer hiPSC-CMs on soft and stiff PDMS substrates, PDMS surface treatment with PD in combination with ECM proteins was employed. Contractility analysis at various time frames, extending up to 90 days post replating, revealed that cells exhibited optimal functionality after 22 days of culturing on soft micropatterned substrates with Center to Center (C-C) distance of 20 mm compared to the other dimensions. Furthermore, the results of electrophysiological analysis in hiPSC-CMs cultured on soft micropatterned PDMS substrate showed faster rise time and decay of Ca^2+^ transients when shifting from 7 to 22 days. While immature hiPSC-CMs cultured on stiff micropatterned substrate exhibited the highest sarcomere shortening and prolonged Ca^2+^ transient duration, mature hiPSC-CMs demonstrated maximal contractility and Ca^2+^ transient on a soft micropatterned substrate. Additionally, micropatterned substrate with Center to Center (C-C) distance of 20 mm shows the highest nucleus aspect ratio, enhanced mitochondria density, and the most uniform sarcomere orientation. These biologically compatible substrates pave the way for disease-in-a-dish studies, representing the pathogenic phenotype of a disease, and offering novel insights for testing therapeutic interventions.

## 2. MATERIAL AND METHODS

### 2a. Substrate fabrication

The novel fabrication process for the soft micropatterned substrate used in this study includes three main steps: 1-stiff PDMS mold fabrication, 2-sacrificial layer formation, and 3-soft PDMS casting and demolding (Figure 1). Firstly, Kapton tape (LINQTAPE PIT0.5S-UT and PIT1S Series Kapton polyimide film tape) was affixed to the glass slide. Following this, grooves were created on the Kapton tape utilizing a commercial laser machining system (QuiLaze 50ST2, New Wave Research, USA) with an UV light (355 nm λ) and 50X NUV microscope lens. The parameters of the UV laser, such as speed, power, and frequency, can be adjusted to modify the spacing, depth, and width of the grooves. In our application, the UV laser specifications were adjusted to ensure a uniform groove depth, allowing the laser to cut through the Kapton tape and create an intact pattern on the glass slide. Laser cutting speed and power have been tested on the effect of width and depth of the groove formation. Once the grooves were established on the Kapton tape, a PDMS (Sylgard 184, Dow Corning) base with a curing agent-to-base ratio of 10:1 was poured onto the grooves to create a PDMS mold as shown in Figure 1A. However, the PDMS mold may degrade after a few demolding processes. Therefore, a permanent mold, utilizing Liquid urethane polymer (Smooth-On Smooth-Cast 305), was employed to create a long-lasting mold. Subsequently, PDMS molds were fabricated from the permanent mold after a couple of uses of the PDMS mold, ensuring the integrity of groove patterns in PDMS mold [7].

**Figure 1.**
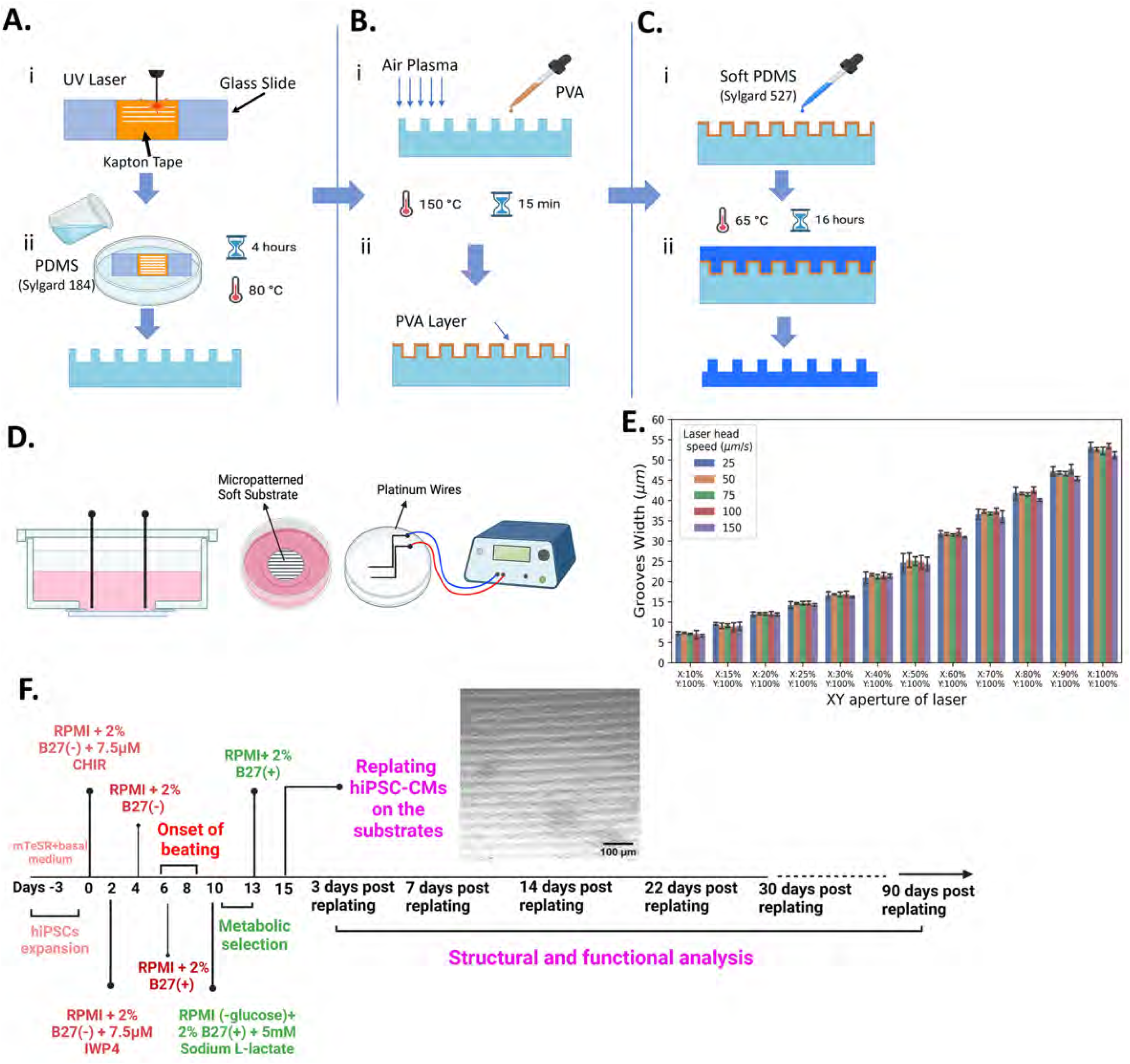
The micropatterned PDMS substrate fabrication process. A) Mold fabrication: A UV laser was used to etch the Kapton tape attached to the glass slide (i) Sylgard 184 with a mass ratio of 10:1 was used to make the mold (ii). B) PVA Sacrificial Layer: Air plasma treatment was applied to enhance the wettability of PDMS mold (i). PVA was poured onto the substrate and cured at 150°C for 15 mins (ii). C) Soft PDMS fabrication: The soft PDMS was fabricated by using Sylgard 527 with a mass ratio of 1:1, and it is poured onto the PDMS mold coated with PVA (i). The soft PDMS was cured in the oven for about 16 hours at 65°C. D) The culture dish is equipped with platinum electrodes within a closed chamber environment, connected to an electrical stimulator for pacing the cells at various frequencies. E) The UV Laser specification for creating patterns with different width on a Kapton tape. F) A timeline for hiPSCs differentiation, purification, and maturation into cardiomyocytes. hiPSC-CMs were replated on glass and PDMS substrates at day 15 post-differentiation. Structural and functional analysis began three days after cardiomyocyte replating and continued until day 90.

Secondly, polyvinyl alcohol (PVA) was employed as a sacrificial layer to facilitate the detachment of soft PDMS from the PDMS mold (Figure 1B). Dissolving PVA in water allows for easy removal of the soft PDMS from the PDMS mold containing grooves. To achieve a uniform PVA layer on the PDMS mold, the first step involved activating the PDMS mold surface through air plasma surface treatment for approximately 2 minutes at a chamber pressure of 700 mTorr. Following this, a specified concentration of PVA was applied to the PDMS mold through the spin coating and then cured on a hot plate at 150 °C for about 15 minutes (Figure 1B).

Finally, to create a soft PDMS, a blend of two commercially available PDMS types, Sylgard 527 (Dow Corning) by mixing parts A and B in a mass ratio of 1:1 and Sylgard 184 (Dow Corning) in a mass ratio of 10:1 (base/curing agent), was utilized (Figure 1C). This PDMS mixture provides the flexibility to adjust the substrate stiffness within a range from 2 kPa to 1.72 MPa, independent of changing other material properties such as surface roughness, surface energy, and the capacity to functionalize the surface through protein absorption [39]. The soft PDMS was coated onto the PDMS mold, which was previously covered by a layer of PVA serving as a sacrificial layer. Subsequently, the soft PDMS underwent a degassing process in a vacuum chamber for approximately 1 hour to eliminate bubbles. The soft PDMS was cured in a preheated gravity convection oven (Yamato DX 300, Integrated Services, Palisades Park, NJ, USA) at 65°C for about 16 hours. After curing the soft PDMS, air plasma surface treatment was applied to activate both the soft PDMS and glass coverslips (25 mm, No. 0, MatTek Corporation) surfaces. Subsequently, the coverslip was attached to the soft PDMS, and the entire sample was immersed in an ultrasonic cleaner device filled with water for approximately 30 minutes at 40°C. This facilitated the easy removal of the soft PDMS by dissolving the PVA layer in water. (Figure 1C).

To measure the depth of the grooves and assess surface topography, a Wyko NT1100 Optical Profiling System was utilized (Figures 2A and 2B). Furthermore, the thickness of the microgrooved PDMS substrate was determined using 0.2 μm fluorescent beads (FluoSpheres™ Carboxylate-Modified Microspheres, Invitrogen by Thermo Fisher Scientific) embedded into the PDMS substrate, and subsequent measurements were conducted using fluorescent microscopy (Figure 2C).

**Figure 2.**
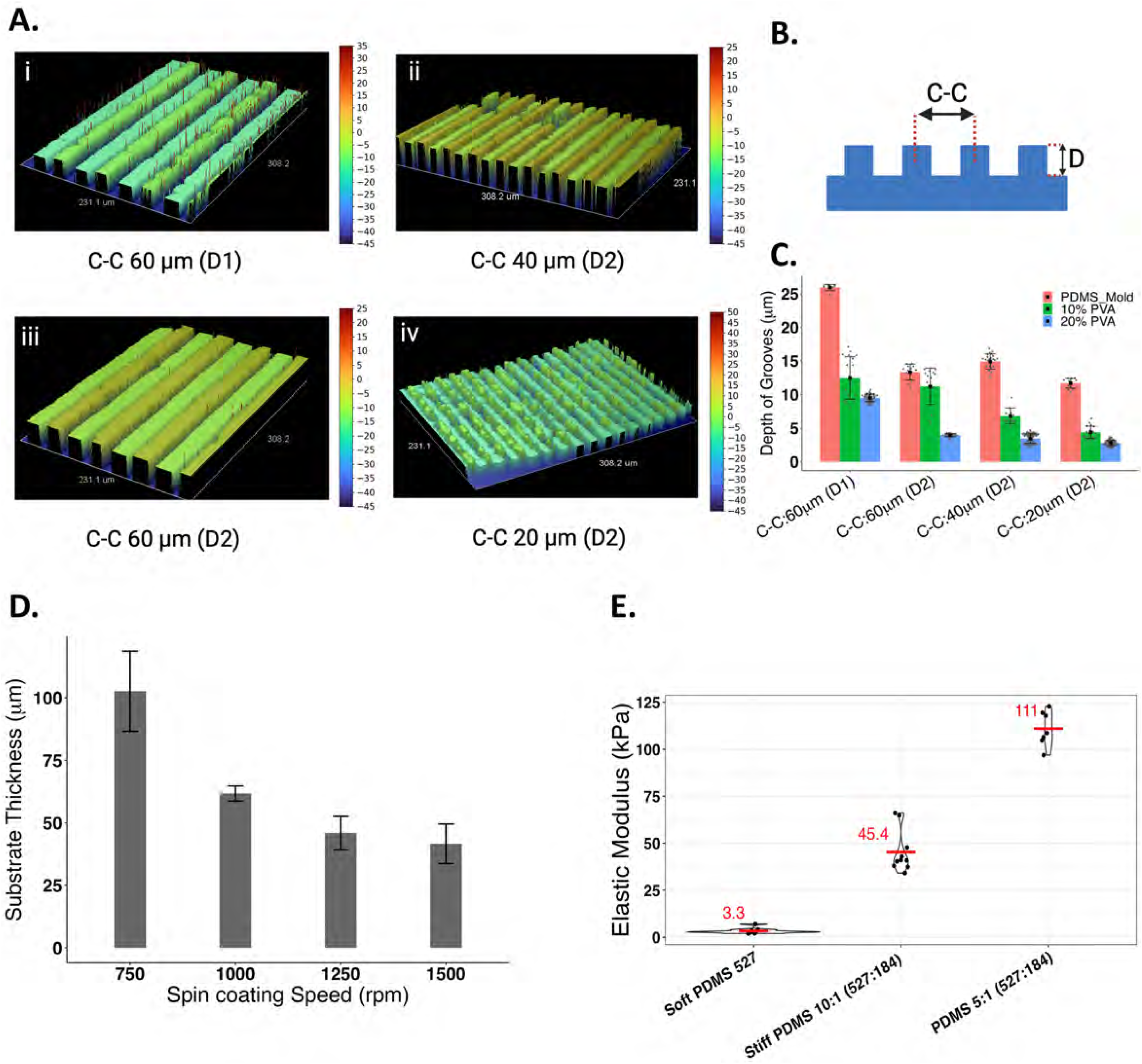
Surface topographical features and specifications of soft micropatterned PDMS substrate characterized by Wyko optical profiler. A) C-C of 60 µm (D1): grooves center to center distance of 60 µm with a depth range of 8 µm to 10 µm (i), C-C of 40 um: center to center distance of 40 µm with the depth range of 2.5 µm to 5 µm (ii), C-C of 60 µm (D2): center to center distance of 60 µm with the depth range of 2.5 µm to 5 µm (iii), C-C of 20 um: center to center distance of 20 µm with the depth range of 2.5 µm to 5 µm (iv). B) Dimensions of the micropatterned substrate. C) The depth of the grooves in the PDMS mold and soft PDMS using 10% PVA (green) and 20% PVA (blue) as a sacrificial layer. D) The thickness of the soft PDMS substrate at different spin coating speeds. E) Stiffness of PDMS with three different formulations using one-term Ogden model.

### 2b. Mechanical characterization of substrate

A uniaxial test was performed using an Instron 5969 Universal Testing System (Instron Corp.) to characterize stiffness of the substrate. Standard dogbone-shaped molds (ASTM D-412) were laser-cut from polymethyl methacrylate (PMMA). To fabricate intact PDMS samples for uniaxial testing, a PVA layer was employed as a sacrificial layer. To fabricate PDMS substrates with different stiffnesses, three sample groups with varying mass ratios of Sylgard 184 to Sylgard 527 were examined, including a ratio of 1:10 (number of samples, n=10), a ratio of 1:5 (n=7), and samples with only Sylgard 527 (n=11). All samples were derived from at least three different preparations. The stretching rate during testing was set at 20 mm min^-1^.

### 2c. Water contact angle

Contact angle measurements using water droplets were conducted to evaluate the surface hydrophilicity of soft micropatterned and unpatterned PDMS substrates with and without PDMS surface treatment. In this study, we exclusively utilized a soft PDMS formulation to measure the water contact angle with and without protein coating. To measure the water contact angle, soft PDMS was poured on a 25 mm glass coverslip. Three different samples from at least two separate preparations were used for each coating condition. The contact angle was assessed using a custom-made setup. A droplet of distilled water (1 µL) was placed on the surface of the substrate, and the average of the contact angles on the left and right sides of the droplet was measured utilizing the Image J Water Angle software plugins (Figure 3)[40].

**Figure 3.**
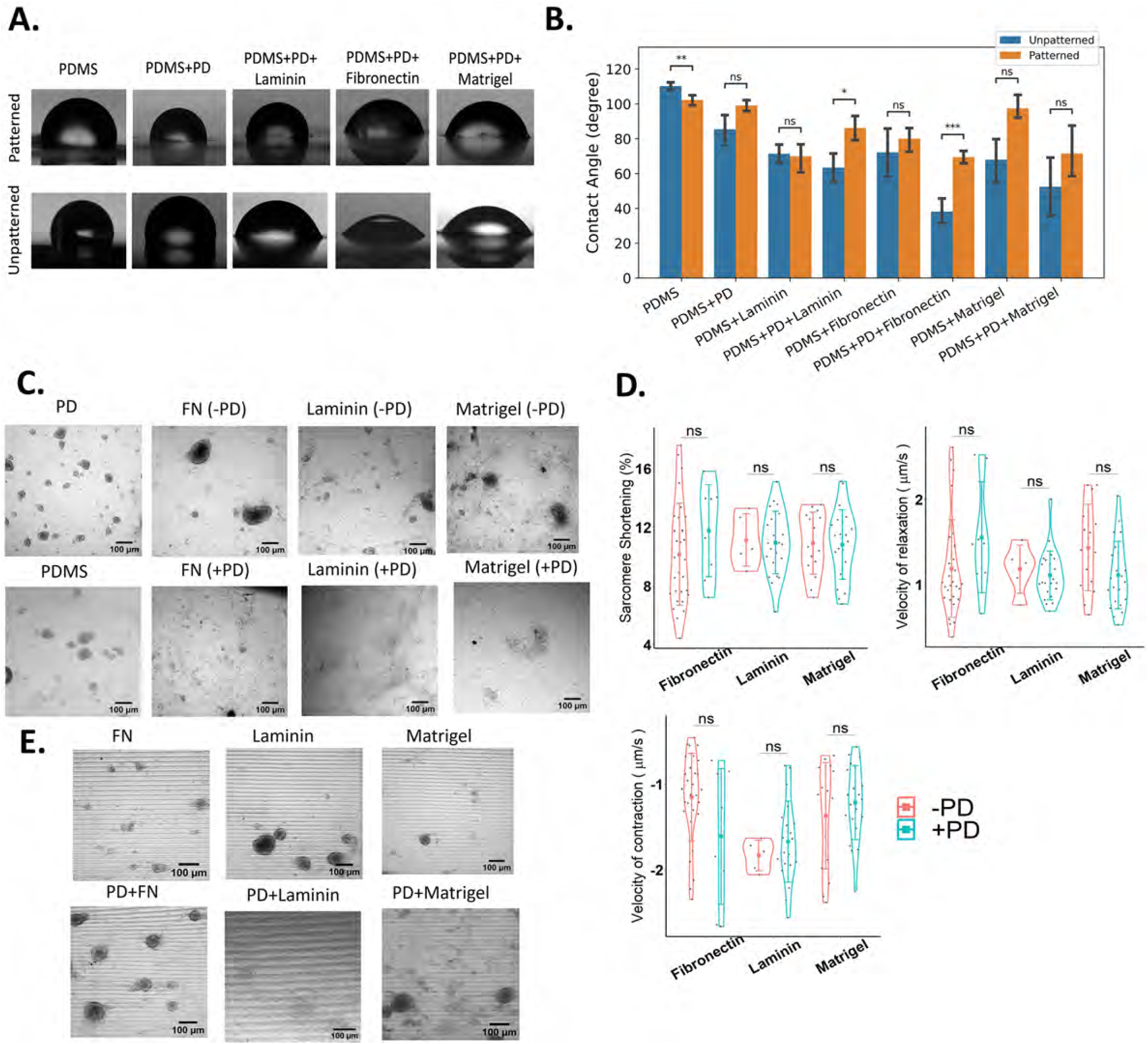
Surface hydrophobicity and surface modification analysis. A) Representative images of water droplet formation on patterned and unpatterned soft PDMS with different surface coatings. B) Quantification of water contact angle for patterned (Orange) and unpatterned (Blue) non-coated PDMS, PD coated PDMS, ECM (Laminin, Fibronectin, Matrigel) coated PDMS, and PD/ECM coated PDMS substrates. C) The attachment of hiPSC-CMs cultured on soft unpatterned non coated PDMS, PD coated PDMS, ECM (Laminin, Fibronectin, Matrigel) coated PDMS, and PD/ECM coated PDMS substrates. The bright field images were captured after 1 week of replating the cells onto the soft PDMS substrates using Nikon Ti2 microscope. D) The percentage of sarcomere shortening, contraction velocity, and relaxation velocity of hiPSC-CMs cultured on unpatterned soft ECM coated (Orange) and PD/ECM coated (Blue) PDMS substrate. E) The effect of patterning on cell attachment on the soft ECM (Laminin, Fibronectin, Matrigel) coated PDMS and PD/ECM coated PDMS substrates. The bright field images were captured after 1 week of replating the cells onto the soft micropatterned PDMS substrates using Nikon Ti2 microscope. ns p-value>0.05, * p-value< 0.05, ** p-value <0.005, *** p-value<0.0005, **** p-value< 0.00005.

### 2d. hiPSC differentiation to CMs

To differentiate hiPSCs into cardiomyocytes in following our previously published protocol [41][42][24], hiPSCs were seeded on a six-well plate coated with Matrigel (0.5 mg/six-well plate, dissolved in DMEM/F-12 medium) in mTeSR Plus medium (StemCell Technologies, Vancouver, BC, Canada). Once reaching 60% confluency, the mTeSR medium was replaced with Roswell Park Memorial Institute Medium (RPMI) 1640 supplemented with 2% B27 (-insulin) and 7.5 μM CHIR99021 for 48 hours. Subsequently, the medium was changed to RPMI supplemented with 2% B27 (-insulin) and 7.5 μM IWP4. After 2 days, the media were further changed to RPMI and 2% B27 (-insulin) for 48 hours, followed by another change to RPMI and 2% B27 (+insulin). Metabolic selection was initiated once spontaneous beating was observed around days 7–10 to ensure a purification level of approximately 95%. Given that cardiomyocytes can utilize lactate as an alternative energy source, metabolic selection was performed cells were replated on both unpatterned and micropatterned PDMS substrates, as well as glass-bottom dishes (No 1.5 cover glass, MatTek Corporation P35G-1.5-14-C) (Figure 1F).

### 2e. PDMS surface treatment

Both patterned and unpatterned PDMS substrates were first coated with a 0.01% dopamine solution in Tris-HCl buffer (pH 8.5) overnight [25][26]. Subsequently, the PD-coated PDMS substrates were double coated with laminin, Matrigel, and fibronectin at a concentration of 50 μg/ml. To find the optimal combination of PD and ECM proteins, hiPSCs were differentiated into cardiomyocytes (CMs) on tissue culture dishes, following the methodology outlined in the previous section. At day 15 of differentiation, the hiPSC-CMs were replated on both micropatterned and unpatterned ECM-coated and PD/ECM-coated PDMS substrates. To evaluate cell attachment after 1 week of replating, bright field images were captured using a Nikon confocal spinning disc Ti2 microscope (Figure 3C and D). Next, the impact of PD coating on the function of hiPSC-CMs was investigated. To achieve this, cell contractility was measured only on unpatterned ECM-coated and PD/ECM-coated PDMS substrates since cells detached on ECM-coated micropatterned substrates after a couple of days. The effect of PD coating on hiPSC-CM contractility was assessed 1 week after replating using SarcTrack (Figure 3B) [43].

### 2f. Live cell imaging

In this study, Allen Institute hiPSCs (AICS-0075) were utilized, in which the α-actinin2 which is expressed in the Z-line of sarcomeres, was genetically tagged with green fluorescent protein (GFP). The nuclei of hiPSC-CMs were stained with 0.8 µM Hoechst (Thermo Fisher). Mitochondria staining was performed using MitoTracker Deep Red (Thermo Fisher), which was diluted in the cell’s media (RPMI without phenol red/B27+) at a ratio of 1:4000. Following a 15-minute incubation of cells in a 37°C incubator with Mitotracker and Hoechst, the cells were washed with D-PBS, and the media were replaced with RPMI (-phenol) and 2% B27 (+ insulin). Live cell imaging was conducted using the Nikon Ti2 confocal spinning disc microscope with a 60x water immersion objective and the Leica confocal SP8 with a 40x objective.

### 2g. Orientational Order Parameter (OOP), nucleus aspect ratio, and mitochondria density

The Orientational Order Parameter (OOP) is a well-established metric used to quantify alignment in biological systems, ranging from 0 to 1. A score of 0 indicates complete isotropic orientation, while a score of 1 denotes perfect anisotropic orientation [44][29]. In our study, the OOP was employed to characterize the orientation of Z-lines in hiPSC-CMs cultured on unpatterned and micropatterned soft and stiff PDMS substrates. In our OOP analysis, the spinning disk confocal images (60X water objective) and Leica SP8 confocal microscopy images (oil objectives 40X and 63X) were utilized to capture the GFP tagged Z-line of the sarcomeres. The GTFiber package in MATLAB, recommended by Persson et al. [45], was employed for Z-line detection and OOP determination in hiPSC-CMs. The image processing workflow of GTFiber included diffusion filtering, top hat filtering, thresholding, cleaning, and skeletonization processes to obtain the skeleton and subsequent orientation map of Z-lines in hiPSC-CMs.

After determining the appropriate orientation map for each image, fibres perpendicular to the Z-lines were removed using a custom pipeline in CellProfiler [46]. The output of the CellProfiler consists of images displaying the Z-lines of the hiPSC-CMs, which were then utilized to calculate the OOP using the GTFiber package. Throughout the image analysis, the grid step and frame step were chosen based on the image size and kept consistent for all images of the same size (see Supplementary Figure S2 and S3). The nucleus aspect ratio was determined with a custom pipeline in CellProfiler. The Minimum Cross-Entropy method was used to threshold the nuclei in the Hoesch channel of the fluorescent images. The nuclei were identified using the IdentifyPrimaryObjects module of CellProfiler.

### 2h. Functional analysis of hiPSC-CMs

**I**n this study, SarcTrack [43]was utilized for four specific purposes to: 1) assess the impact of PD on hiPSC-CMs contractility, 2) explore the influence of patterning on resting (diastole) and contracting (systole) sarcomere length at intervals of 3, 7, 16, 22, 30, and 90 days while cultured on both unpatterned and micropatterned substrates with three different groove dimensions of C-C 20 µm, 40 µm, and 60 µm, along with D2 depth (2-5 μm), 3) identify the optimal time frame wherein the resting sarcomere length reaches its maximum value, ideally close to 2.0 -2.2 µm, indicating maturation, and 4) examine the effect of substrate stiffness on hiPSC-CMs contractile parameters.

The SarcTrack algorithm fitted each sarcomere to a double wavelet, employing six definable parameters: 1) distance between wavelet peaks as a function of time, 2) constraint length of each wavelet, 3) width of each individual wavelet, 4) number of angles that wavelet test to find the best fit to the z-disc, 5) distance between grid points to avoid refitting same sarcomere, and 6) the neighborhood around each grid where the best fit for the wavelet pair can be determined. Images were captured at a rate of 100 frames per second (fps) over a 5-second duration, resulting in a total of 500 frames. Each frame underwent analysis using the SarcTrack algorithm to quantify the variation in sarcomere length over time. The images were acquired using a spinning disc confocal Nikon Ti2 microscope at a frame rate of 100 frames per second (fps), with hiPSC-CM monolayers being stimulated at 1 Hz and maintained at 37°C. To image fluorescently labeled sarcomeres, the α-actinin 2-GFP tagged hiPSC-CMs (Allen cell line (AICS-0075)) were used.

Images were acquired at specific intervals of 3, 7, 14, 22, 30, and 90 days after replating cells onto the patterned and unpatterned soft and stiff PDMS substrates as well as glass dishes. Subsequently, hiPSC-CMs were stimulated at 1 Hz and 5 second videos were captured at 100 fps with 60x water immersion objective and ROI of 1626× 1160 pixels. Each video was then split into 6 videos, each containing approximately 200 sarcomeres. A minimum of 40 videos from three distinct replicates were analyzed for each condition, resulting in a comprehensive analysis of at least 8000 sarcomeres. Subsequently, multiple videos were processed in parallel by setting appropriate SarcTrack parameters. To efficiently handle more than 300 videos simultaneously, the files were uploaded to the computational cluster at the Digital Research Alliance of Canada. This advanced research computing system allows the specification of varying numbers of CPUs, effectively reducing computational costs. In this study, 16 CPUs were designated to process over 300 videos in approximately 4 hours. The measured contractility parameters encompassed the resting sarcomere length (peak of diastole), contracting sarcomere length (peak of systole), percentage of sarcomere shortening, maximum velocity of contraction, and maximum velocity of relaxation of hiPSC-CMs.

### 2i. High speed optical mapping (OM) for Ca^2+^ transient measurements

The 2D monolayer of hiPSC-CMs cultured on soft and stiff PDMS substrates were incubated with 5 μM of the Ca^2+^ sensitive dye Cal630 (AAT Bioquest), diluted in the cell’s media, for 1 hour in a 37°C incubator. Subsequently, the cells were washed with DPBS (without Mg^2+^ and Ca^2+^, STEMCELL Technologies), and the media was replaced with RPMI (without phenol red, Gibco, ThermoFisher Scientific) and 2% B27 (+insulin). The culture dish was equipped with a platinum electrode in a closed chamber environment, maintaining a constant temperature of 37°C and 5% CO2 during data acquisition. For macroscopic optical mapping, the hiPSC-CMs were excited by a 580 nm LED light source (Lumencore) and the emission signal was captured with a high-speed camera (Scimedia MiCAM05) at 277 fps. Data analysis was conducted using our custom MATLAB code, following some of the protocols outlined in a previous [47].

To analyze the calcium transient data, the Ca^2+^ signals were acquired from the optical mapping setup, which is based on the fluorescent intensity of Cal630 within the region of interest (ROI). This intensity is represented as the difference between the background fluorescent intensity and the intensity of Cal630 in each ROI. Prior to analysis, a moving average filters were applied to the raw data and window sizes of 20 to 40 ms were selected to smooth the raw fluorescent intensities of Ca^2+^ traces.

For the identification of the onset of each Ca^2+^ transient signal, the central difference method was used to compute the derivative of the filtered signals. The peak of the derivative of the signals served as the marker for the onset of the calcium transient. To determine the onset of each single Ca^2+^ transient, an offset equivalent to 10% of the cycle length were subtracted from the peak of the calcium transient signal. For instance, this offset was determined to be 100 ms during 1Hz pacing frequency. Consequently, individual traces chosen for analysis fell between two successive offset points.

Subsequently, a baseline intensity was determined to measure all necessary temporal parameters for analyzing the calcium transient. The baseline was determined by averaging the points corresponding to the last temporal window of 20% of the cycle length. For example, a window size of 200 ms was selected for a 1 Hz pacing frequency. The peak of the trace was identified as the maximum intensity value corresponding to the calcium transient trace.

To address potential variations in baseline intensity, Psaras et al. proposed a more robust baseline calculation [47]. Given that a fluorescent trace might exhibit a lower baseline at the end compared to the beginning, the robust baseline was defined as the baseline plus 0.03 times the magnitude. Here, the magnitude represents the absolute difference between the baseline and the peak of the fluorescent trace. This robust baseline was used to compute all temporal parameters through linear interpolation of time at intensity values corresponding to 0% and 50% of the magnitude. All the temporal parameters used in the calcium transient analysis showed in Figure S 5.

### 2j. Statistical analysis

Statistical analyses were performed using R version 4.3.1. All data were reported as mean ± SEM. To evaluate the significance level among multiple independent variables, a one-way or two-way ANOVA test was employed. To assess the significance level between pairs of independent variables, a pairwise comparison t-test with False Discovery Rate (FDR) adjustment for p-values was employed, and statistical significance was defined as a p-value less than 0.05.

## 3. RESULTS

### 3a. Optimizing fabrication parameters

In this study, we utilized PDMS to create a physiologically relevant environment for hiPSC-CM maturation, considering substrate stiffness and patterning. A UV laser etching method was employed to generate patterns on Kapton tape. Subsequently, the mold casting technique, incorporating a sacrificial layer, was used to transfer these patterns onto the soft PDMS substrate. the entire fabrication process can be conducted in a standard laboratory setting without requiring a cleanroom (Figure 1).

In our investigation, the laser specifications played a crucial role in achieving precise and smooth patterning on the Kapton tape. Hence, Further experiments were conducted to explore the impact of varying power levels and X-Y stage speeds using the rectangular laser spot with laser pulses at 50 Hz (Figure 1E). Experimental results demonstrate that using a rectangular laser spot with a longer length along the grooves’ direction, at a speed of 75 μm/s, not only produces intact microgrooves but also does so efficiently in terms of fabrication time. Moreover, the minimum consistent resolution for center-to-center (C-C) spacing of grooves was 20 μm, resulting in making microgrooves 10 μm wide. The optimized laser parameters for creating grooves with various C-C spacing on Kapton tape are detailed in Table 1.

**Table 1:** Laser etching specifications in different power and laser speed.

| C-C Spacing (μm) | Power (Laser Spot) Specifications | Speed (μm/s) | Laser pulses | Number of passes | Energy Range | Kapton Tape |
| --- | --- | --- | --- | --- | --- | --- |
| 20 | 100% energy, High, X (15%), Y(100%) | 75 | 50 Hz, Continuous | 1 | High | PIT0.5S-UT |
| 40 | 100% energy, High, X(30%), Y(100%) | 75 | 50 Hz, Continuous | 1 | High | PIT0.5S-UT |
| 60 | 100% energy, High, X(50%), Y(100%) | 75 | 50 Hz, Continuous | 1 | High | PIT0.5S-UT |
| 60 | 100% energy, High, X(50%), Y(100%) | 75 | 50 Hz, Continuous | 2 | High | PIT1S |

To investigate the effect of PVA concentration and spin coating speed on the integrity of microgrooves, the PDMS mold was coated with PVA at different concentrations including 1 wt%, 10 wt%, and 20 wt%. For PVA 1 wt%, it was difficult to peel off an intact soft PDMS, and the grooves could not be transferred from the PDMS mold to the soft PDMS. Hence, samples with 20% and 10% PVA concentration were prepared, and the depth of the grooves was measured by using The Wyko Optical Profiler system. The results in Figure 2C indicate that a 20% PVA concentration of the sacrificial layer enables the fabrication of more uniform and intact groove patterns with lower depth variation compared to the 10 % PVA concentration. Furthermore, through experimentation with various spinner speeds for coating the PVA sacrificial layer, ranging from 500 to 5000 rpm for 30 seconds, it was observed that a spinner speed of 1500 rpm is the optimal choice for achieving consistent microgrooves compared to other speeds. After applying the PVA as sacrificial layer to the PDMS, the soft PDMS was poured onto the sacrificial layer using a spinner. Following the fabrication process, micropatterned substrates were created with center to center of 60 µm and variable depths of D1 (ranging from 8 to 10 µm) and D2 (ranging from 2.5 to 5 µm) (Figure 2A, i, iii). Subsequently, the depth of grooves was maintained at D2 while the center to center of grooves was reduced to 40 µm and 20 µm (Figure 2A, ii, iv).

Considering the limited working distance of the 60X water immersion objective, approximately 200 μm, and the thickness of the glass coverslips, ranging from 80 to 130 μm, the substrate thickness was controlled to be less than 100 μm. Various spinner speeds were tested for 30 seconds to achieve a specific substrate thickness as illustrated in Figure 2D. It was observed that a spinner speed of 1000 rpm resulted in a uniform thickness with less variation. Hence, all soft PDMS substrates were fabricated with consistent specifications, including curing time and spinner speed in each stage, to ensure a uniform thickness range. Additionally, it is crucial to avoid using a very thin layer of soft PDMS because the substrate stiffness changes at lower thickness levels, leading cells to sense the stiff glass at the bottom. Hence, consistency in substrate thickness is vital for reliable and meaningful results in studying the effect of substrate stiffness on hiPSC-CMs maturation and functionality [30][44].

### 3b. Mechanical properties of PDMS substrate

Three hyperelastic constitutive models, including the one-term Ogden model, Neo-Hookian, and Moony-Rivlin models, were employed to characterize the mechanical behavior of PDMS samples with three different formulations by blending Sylgard 527 and Sylgard 184 [48], [49]. Assuming incompressibility, the axial tension λ = 1 + |ϵ| generates an elastic Cauchy stress *T*^*e*^, which is detailed in Table 2 for three different hyperelastic models. According to the Root Mean Square Error (RMSE) (See Supplementary Figure S1), the one-term Ogden model exhibits a better fit compared to the other models, consistent with previous studies [48], [49]. The elastic modulus based on the one-term Ogden model is given by [50]:

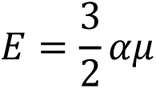

**Table 2:**
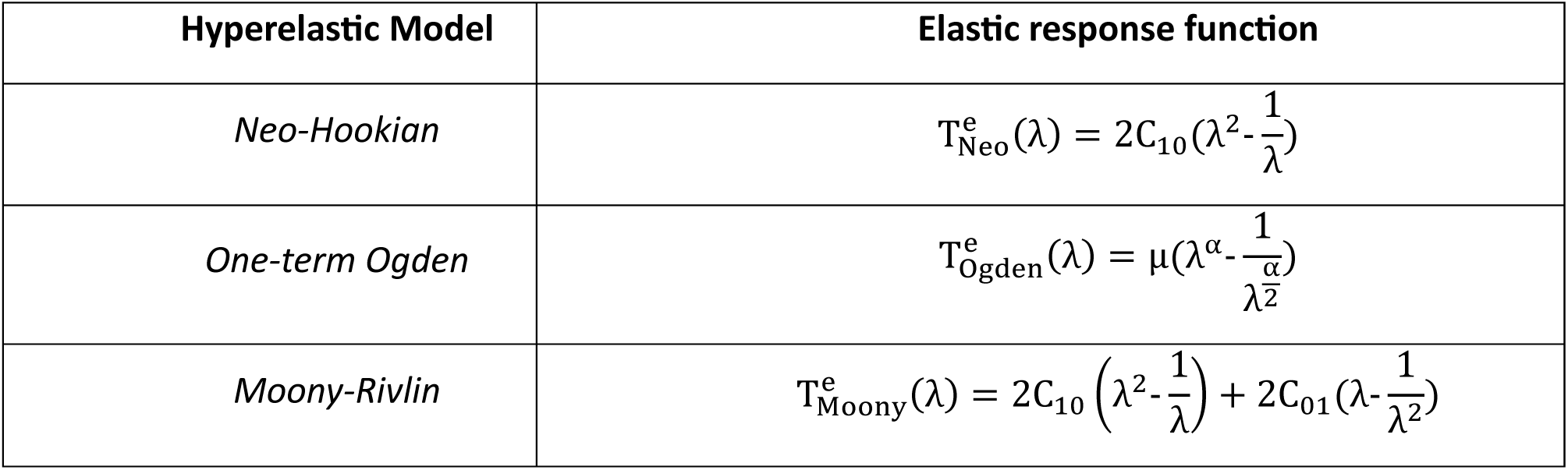
The hyperelastic models used to characterize the substrate.

Figure 2E represents the elastic modulus for three different PDMS sample groups, modelled using the one-term Ogden model.

### 3c. PDMS surface modification with PD

The contact angle of water droplets was measured for the cells cultured on both unpatterned and micropatterned soft PDMS substrate coated with PD/ECM, as well as substrates coated with only ECM proteins as illustrated in Figure 3A and B. The PD/fibronectin coating exhibited the lowest contact angle on both patterned and unpatterned substrates, whereas the highest angle was observed for PDMS and PD coating without ECM proteins (Figure 3B). Coating substrates with PD and ECM proteins in a two-step process significantly reduces the water contact angle compared to non-PD-treated substrates (Figure 3A and B).

To determine the effect of surface topography on cell adhesion, hiPSC-CMs were cultured on both patterned and unpatterned PDMS substrates coated with only ECM and PD/ECM proteins. The area of cell attachment was monitored after a week using the Nikon Ti2 microscope. As shown in Figure 3C, on unpatterned soft PDMS substrates coated with only PD and non-coated PDMS substrates, cells exhibited clumping initially and complete detachment after a week. However, no cell detachment was observed for hiPSC-CMs cultured on the dual coated PD/ECM unpatterned PDMS substrates. Among the ECM-coated group, the unpatterned substrate coated with laminin exhibited the lowest cell attachment (Figure 3C). In contrast, the PD/laminin-coated substrate showed the largest cell area attachment (Figure 3C). Cell area assessment was also conducted for hiPSC-CMs cultured on micropatterned PDMS substrates, including two conditions of PD-treated and non-PD-treated. As illustrated in Figure 3E, following one week of replating cells on the micropatterned substrate, only cells on PD/Laminin-coated substrates remained attached and exhibited the largest cell area.

Subsequently, the influence of PD coating on the function of hiPSC-CMs was examined, given previous findings suggesting that PD could impact the surface stiffness of PDMS [25]. To investigate this, cell contractility was measured exclusively on unpatterned ECM-coated and PD/ECM-coated PDMS substrates, as cells detached on ECM-coated micropatterned substrates after a couple of days. The impact of PD coating on hiPSC-CM contractility was evaluated one week after replating the cells on unpatterned PDMS substrates using SarcTrack. As depicted in Figure 3D, the percentage of sarcomere shortening, contraction velocity, and relaxation velocity of hiPSC-CMs remained unchanged on unpatterned substrates coated with ECM compared to those coated with PD/ECM.

### 3d. The effect of patterning on the orientation of sarcomeres and nuclei

To examine the impact of pattern size on the orientation of hiPSC-CMs, cells were replated on unpatterned and patterned substrates with varying C-C groove distancing of 20 µm, 40 µm and 60 µm. First, the OOP were measured for the cells on an unpatterned soft PDMS substrate. As shown in Figure 4A, a disorganization in the sarcomere structure with an OOP of about 0.2 was observed. Furthermore, the pronounced variability in the OOP distribution confirms the unpatterned substrate’s inability to effectively align the hiPSC-CMs in the direction of their contraction, thereby hindering the potential for synchronized beating. Next, to investigate the effect of groove depth on sarcomere orientation, cells were replated on a micropatterned substrate with a C-C of 60 µm and varying depths of D1 and D2. Although hiPSC-CMs fully occupied the 8 to 10 µm-deep grooves on the patterned substrate, no alignment was observed along the pattern direction (Figure 4A). The OOP distribution exhibited lower variability in comparison to the unpatterned substrate; however, the OOP value did not show a significant increase when compared to cells on an unpatterned substrate (Figure 4B and C). Due to the considerable depth of the grooves, a multilayer of cardiomyocytes becomes trapped within them, hindering the cells from effectively sensing and responding to the underlying patterns. While no alignment was evident for hiPSC-CMs on the patterned substrate with a depth of D1, a notable increase in both sarcomere alignment and OOP was observed for cells on the patterned substrate with depth of D2, as illustrated in Figure 4A, B, and C. The increased uniformity in the OOP distribution for cells on substrates with a D2 depth compared to D1, and the relatively high OOP value of approximately 0.5 for D2 deep substrates emphasize the significance of groove depth in influencing the alignment of cardiomyocytes (Figure 4B).

**Figure 4.**
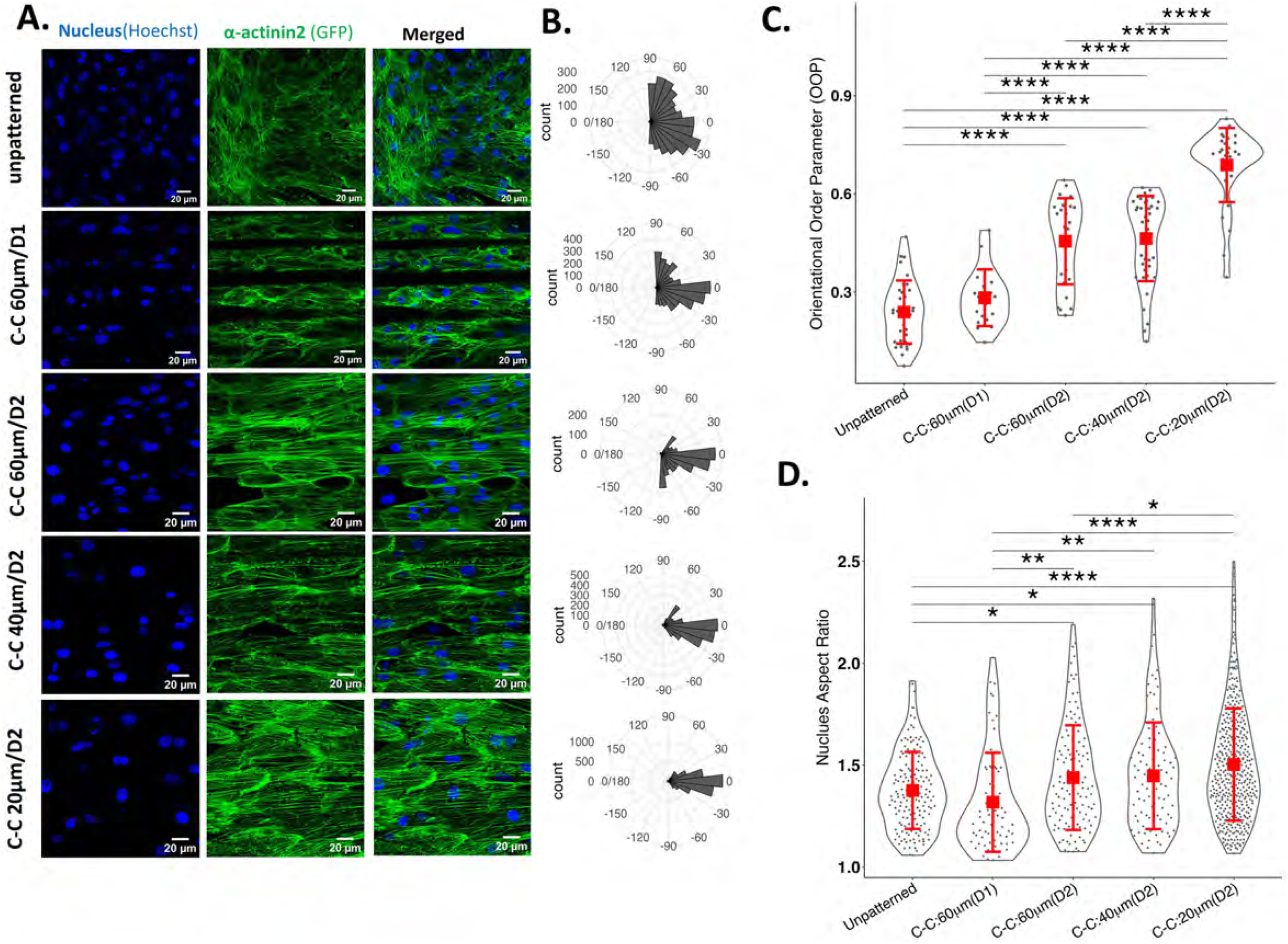
The effect of different pattern sizes on sarcomere orientation, OOP distribution, and nucleus aspect ratio. A) Live cell imaging of sarcomeres (green) and nuclei (blue) of hiPSC-CMs cultured on both unpatterned and micropatterned soft PDMS substrate with varying pattern dimensions: C-C 60 µm/ depth 10 µm, C-C 60 µm /depth 5 µm, C-C 40 µm /depth 5 µm, C-C 20 µm /depth 5 µm. The scale bar is equal to 20 μm. B) Distribution of sarcomere direction with respect to the horizontal orientation of the patterns for both unpatterned and micropatterned soft PDMS substrate with varying pattern dimensions. C) OOP value, and D) Nucleus aspect ratio for hiPSC-CMs cultured on both unpatterned and micropatterned soft PDMS substrate with varying pattern dimensions.

After identifying the optimal depth for hiPSC-CMs alignment, the depth was maintained constant at D2, while the C-C was systematically varied to 40 µm and 20 µm to investigate the optimal C-C distance for achieving the highest OOP and sarcomere organization, indicating mature cardiomyocytes. As shown in Figure 4A and B, the alignment and OOP distribution of hiPSC-CMs on the 40 µm pattern were slightly higher, although not significantly, compared to those on the 60 µm pattern. The OOP value for hiPSC-CMs on 40 µm patterned substrates closely resembled that of the 60 µm C-C and D2 depth, but it was significantly higher than the unpatterned substrate as well as the 60 µm C-C with D1 depth grooves (Figure 4C). In contrast, hiPSC-CMs on the 20 µm C-C patterned substrate with D2 depth displayed the highest alignment, the most uniform OOP distribution, and the highest OOP value of about 0.8 as depicted in Figure 4A, B, and C. Moreover, hiPSC-CMs cultured on this substrate exhibited rhythmic beating aligned with the pattern as demonstrated in Supplementary Video 2. This alignment of cardiomyocytes was significantly greater than all other pattern dimensions as well as the unpatterned substrate.

Moreover, the impact of patterning on nuclear dimensions was studied, as morphological changes in cells have been linked to alterations in nucleus size. As depicted in Figure 4D, the aspect ratio of the nucleus is approximately the same for both unpatterned and patterned substrates with a C-C of 60 µm and a depth of D1. However, when reducing the groove depth from D1 to D2, a significant increase in the nucleus aspect ratio was observed compared to both the D1 deep patterned and unpatterned substrates (Figure 4D). This indicates that the depth of the patterns not only affects sarcomere organization but also influences nucleus orientation, ellipticity, and alignment. When reducing the pattern C-C to 40 µm, no significant changes were observed in the nucleus aspect ratio compared to the 60 µm C-C patterns, while a dramatic increase was noted compared to the unpatterned substrate and C-C of 60 µm with D1 depth pattern. As depicted in Figure 4D, decreasing the pattern C-C to 20 µm significantly increases the nucleus aspect ratio, resembling the adult cardiomyocyte phenotype. Culturing hiPSC-CMs on a micropatterned PDMS substrate dramatically enhances maturation in terms of sarcomere organization and nucleus orientation.

### 3e. Enhancing hiPSC-CM maturation with soft micropatterned PDMS substrate

Since no changes in the OOP and nucleus aspect ratio were observed for micropatterned substrate with C-C of 60 µm and depth of D1 compared to unpatterned substrates (Figure 4), SarcTrack analysis was conducted on patterned substrates with C-C of 20 µm, 40 µm, and 60 µm, all having the same depth of D2 (Figure 5 and Figure S 6). As depicted in Figure S6, after 3 days of culturing the cells on micropatterned and unpatterned substrates, the resting and contracting sarcomere lengths remain unchanged for C-C of 20 µm and 40 µm patterns, while they were significantly lower for C-C of 60 µm and unpatterned substrates. After 7 days, the resting and contracting sarcomere lengths increased for cells cultured on a 60 µm (C-C) substrate, reaching the same sarcomere length value as cells on C-C of 20 µm and 40 µm substrates, which is significantly higher than that observed for the unpatterned substrate (Figure S6). On day 14 after replating, the resting sarcomere length for cells on the unpatterned substrate also increased and reached the same value as the patterned substrate, while the contracting sarcomere length remained significantly lower than that of the patterned substrate (Figure S6). The sarcomere length during contraction and relaxation reaches its highest value after 22 days of culturing hiPSC-CMs on micropatterned substrates with 20 µm (C-C) and depth of D2, compared to all other conditions (Figure 5A1 and A2). Additionally, no significant differences were observed in the resting and contracting sarcomere length for cells on C-C of 40 µm, 60 µm, and unpatterned substrates at day 22 (Figure 5A1 and A2). While no changes in the sarcomere length were detected for cells on 40 µm (C-C) compared to the 20 µm (C-C) pattern at day 30, a significant decrease was observed for cells on 60 µm (C-C) compared to both 20 µm (C-C) and 40 µm (C-C) substrates (Figure S 6). Finally, after 90 days of culturing the cells on the soft PDMS patterned and unpatterned substrates, the resting and contracting sarcomere length for cells on the unpatterned substrate was significantly lower than the micropatterned substrate (Figure S6). As depicted in Figure 5A, hiPSC-CMs on the micropatterned substrate with C-C of 20 µm and depth of D2 exhibited the highest resting and contracting sarcomere length throughout all days of culturing.

**Figure 5.**
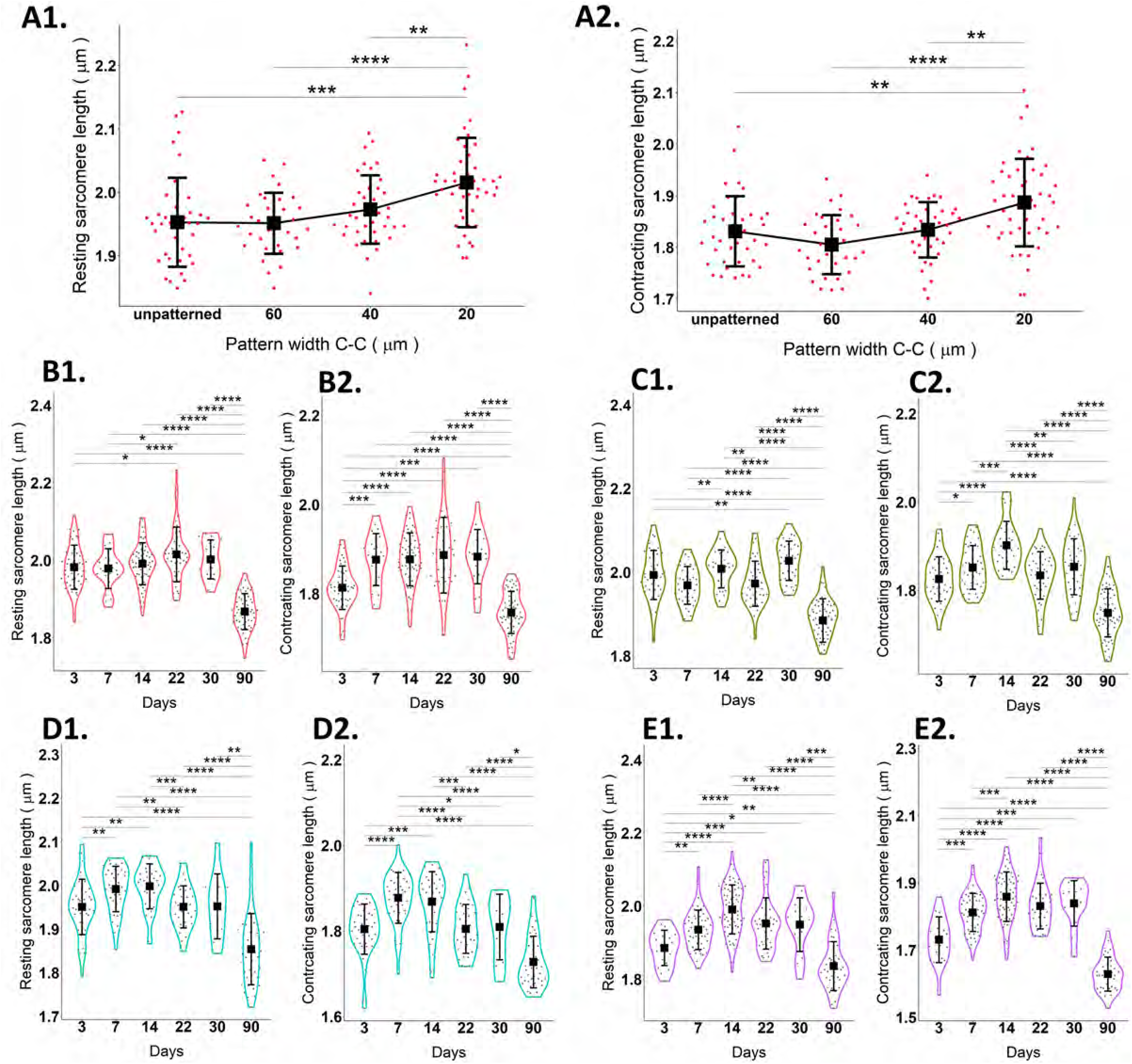
The impact of different pattern sizes and timing on the sarcomere length of hiPSC-CMs. A1) Resting sarcomere length, and A2) contracting sarcomere length of hiPSC-CMs cultured on unpatterned and micropatterned PDMS substrates for 22 days with different pattern dimensions of C-C of 20 µm, 40 µm, and 60 µm with a depth of 5 μm. B1) Resting and B2) contracting sarcomere lengths for hiPSC-CMs cultured on micropatterned substrates with different pattern widths of 20 µm (red), C1 and C2) 40 µm (green), D1 and D2) 60 µm (blue), E1 and E2) unpatterned (purple) substrate at intervals of 3, 7, 14, 22, 30, and 90 days post-replating. A minimum of 40 videos from three different replications were analyzed for each condition utilizing SarcTrack. ns p-value > 0.05, * p-value < 0.05, ** p-value < 0.005, *** p-value < 0.0005, **** p-value < 0.00005.

Next, to investigate the effect of timing on the maturation of hiPSC-CMs and determine the optimum time to achieve the highest sarcomere length, each substrate with a different pattern size was studied individually at various days. As depicted in Figure 5B, for the pattern C-C of 20 µm, the resting sarcomere length at day 22 was significantly higher than at days 3 and 7, and the contracting sarcomere length shows a dramatic increase with the duration extending from 3 to 22 days. After 30 days, the resting and contracting sarcomere length decreased slightly but not significantly compared to days 22, 14, and 7, yet it remained dramatically higher than the length observed on day 3. However, after 90 days, both the resting and contracting sarcomere lengths decreased significantly compared to days 3, 7, 14, 22, and 30. Figure 5C illustrates the effect of timing on the sarcomere length of cells cultured on a 40 µm (C-C) substrate. The highest resting and contracting sarcomere lengths were achieved after 14 days, while the lowest sarcomere length was observed at day 90. Similar results were obtained for 60 µm (C-C) and unpatterned substrates, with the highest resting sarcomere length being less than 2 µm, as depicted in Figure 5D and 5E, respectively. A significant increase in the resting and contracting sarcomere length for cells on the soft unpatterned substrate at day 14 compared to day 3 indicated that not only does pattern affect structural maturation, but timing also plays an important role in the maturation of hiPSC-CMs (Figure 5E). Moreover, a significant reduction in the sarcomere length at day 90 compared to all other days was observed for all pattern dimensions as well as the unpatterned substrates.

### 3f. Electrophysiological studies in a 2D monolayer of hiPSC-CMs cultured on soft micropatterned PDMS substrates

To further determine the optimal time-point for achieving the highest functionality, hiPSC-CMs were cultured on soft micropatterned substrates with a C-C spacing of 20 μm, and the kinetics of Ca^2+^ transients were measured at various time points of 7, 14, 22, and 30 days post-replating. As contractility analysis revealed a significant decrease in sarcomere length at day 90 (Figure 5), this condition was excluded from Ca^2+^ transient measurements.

Initially, a monolayer of hiPSC-CMs was paced at a frequency of 1 Hz, and Ca^2+^ transient kinetics were measured using high-speed optical mapping (OM) (Figure 6). The kinetics of Ca^2+^ transients accelerated from day 7 to days 14-30 post-replating. A significant decrease was observed in the half-rise time (TD50_on) and half-decay time (TD50_off) at day 14 compared to day 7 (Figure 6A and 6B). The half-decay time (TD50_off) was significantly lower at day 22 compared to other days, indicating faster Ca^2+^ reuptake at 50% decay for hiPSC-CMs cultured on soft micropatterned substrates for three weeks, likely due to enhanced SERCA activity or expression as a result of maturation (Figure 6B). Moreover, the Ca^2+^ transient amplitude showed the highest value at day 22, which is significantly higher than all other conditions (Figure 6C). Changes in Ca^2+^ transient kinetics at different post-replating time points are evident from the normalized fluorescent intensity Ca^2+^ traces, with the fastest decay observed for hiPSC-CMs cultured on a soft micropatterned substrate for 22 days (Figure 6D).

**Figure 6.**
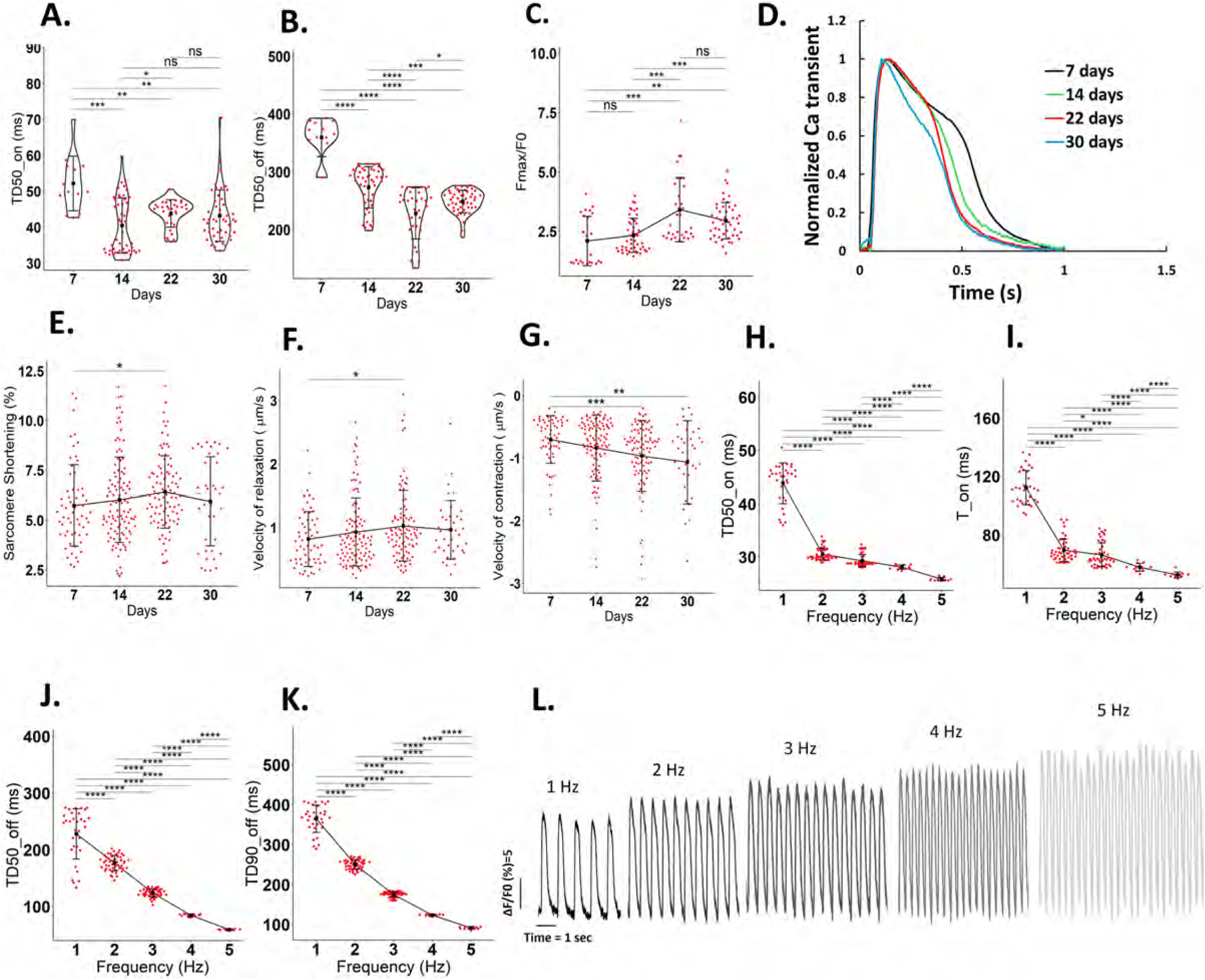
The impact of timing on the Ca2+ transient and contractility of hiPSC-CMs cultured on a soft micropatterned substrate with a C-C distance of 20 μm. A) Half rise time, and B) Half decay time of Ca2+ transient at various days of 7, 14, 22, and 30 post-replating. C) Ca2+ transient amplitude of hiPSC-CMs at various days of 7, 14, 22, and 30 post-replating. D) Normalized Ca2+ transient signal of hiPSC-CMs at various days of 7 (black), 14 (green), 22 (red), and 30 (blue) post-replating. Each Ca2+ transient trace represents the average of at least 120 single traces. E) Percentage of sarcomere shortening, F) Velocity of relaxation, and G) Velocity of contraction of hiPSC-CMs at various days of 7, 14, 20, and 30 post-replating. A minimum of 40 videos from three different replications were analyzed for each condition utilizing SarcTrack. The effect of increasing frequency on H) Half rise time, I) Time to peak, J) Half decay time, and K) Time to 90% decay of hiPSC-CMs at day 22 of replating on the soft micropatterned PDMS substrate. L) Raw Ca2+ transient signals at various frequencies of 1, 2, 3, 4, and 5 Hz. ns p-value > 0.05, * p-value < 0.05, ** p-value < 0.005, *** p-value < 0.0005, **** p-value < 0.00005.

The results of contractility analysis using SarcTrack revealed that the maximum sarcomere shortening, and the fastest contraction and relaxation observed in hiPSC-CMs cultured for 22 days on soft micropatterned PDMS substrate (Figure 6E, F, and G). The sarcomere shortening was significantly increased when shifting from day 7 to day 22 and decreased slightly, but not significantly, from day 22 to day 30 (Figure 6E). Additionally, the velocity of contraction and relaxation significantly increased from day 7 to day 22 and remained relatively constant until day 30 (Figure 6F and G). Therefore, the decreased time to peak, faster decay, enhanced Ca^2+^ transient amplitude, and increased contractility observed from day 7 to days 14-30 all represent the maturation of hiPSC-CMs, with optimal values observed at day 22.

Next, to confirm that three weeks is the optimum time for hiPSC-CM maturation on a soft micropatterned substrate, cells were paced at various frequencies from 1 Hz to 5 Hz (Figure 6H-L). Our data revealed that only hiPSC-CMs cultured on a soft micropatterned substrate for 3 weeks were able to entrain normally to a 5 Hz frequency (Figure 6L). Furthermore, as depicted in Figure 6H-6I, the time to peak, half-rise time, decay time, and half-decay time all decreased with increasing frequency, representing the restitution concept in cardiomyocytes. Additionally, no arrhythmia was observed for hiPSC-CMs upon increasing the frequency from 1 Hz to 5 Hz (Figure 6L).

### 3g. The effect of stiffness on sarcomere and nuclei orientation, mitochondria density, hiPSC-CMs contractility, and cytosolic Ca^2+^ transient

To investigate the effect of stiffness on sarcomere orientation, nucleus dimensions, mitochondria density, and cell contractility, hiPSC-CMs were replated on soft and stiff PDMS matrices with C-C of 20 µm and unpatterned, as well as glass substrates. As depicted in Figure 7A, B, and C, the sarcomere orientation and OOP were similar for cells on glass, soft, and stiff unpatterned substrates. However, cardiomyocytes on both soft and stiff micropatterned substrates exhibited significantly higher OOP and a much more uniform alignment in the direction of the patterns compared to unpatterned substrates (Figure 7A-C).

**Figure 7.**
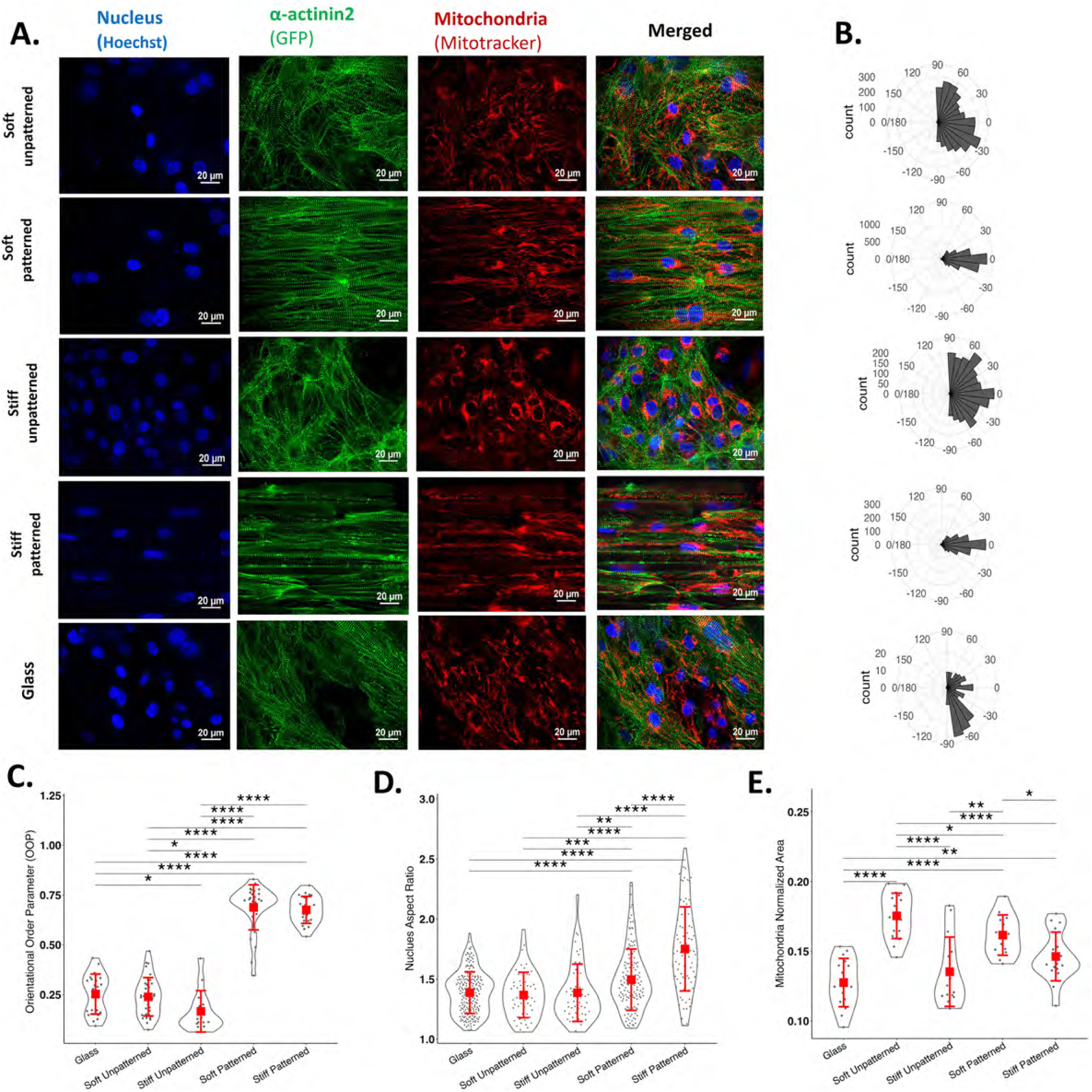
The effect of stiffness on sarcomere orientation, OOP distribution, nucleus aspect ratio, and mitochondria density. A) Live cell imaging of sarcomeres (green), nuclei (blue), and mitochondria (red) of hiPSC-CMs cultured on both soft and stiff unpatterned and micropatterned substrates with a C-C of 20 µm, as well as glass substrate. B) Distribution of sarcomere direction with respect to the horizontal orientation of the patterns for both soft and stiff unpatterned and micropatterned PDMS, as well as glass substrates. C) OOP, D) nucleus aspect ratio, and E) mitochondria density for hiPSC-CMs cultured on both soft and stiff unpatterned and micropatterned substrates with a C-C of 20 µm, as well as glass substrate. The analysis considered a minimum of three replicates, with at least 5 images captured from each replicate. Images were acquired using a confocal spinning disc Nikon Ti2 microscope with a 60x water immersion objective and a field of view (FOV) of 1626 x 1160 pixels. The scale bar is 20 μm. ns p-value > 0.05, * p-value < 0.05, ** p-value < 0.005, *** p-value < 0.0005, **** p-value < 0.00005.

Moreover, the nucleus aspect ratio analysis for cells on glass, unpatterned, and micropatterned substrates was conducted (Figure 7D). The nucleus, being mechanosensitive and mechanoresponsive, plays a critical role in modulating gene expression in response to external stimuli [52][53]. Among various parameters commonly employed in nucleus morphology analysis, this study specifically focused on the aspect ratio, defined as the ratio of the major to minor axis of the nucleus, showing ellipticity of nuclei. This choice was made because this ratio remains relatively constant across different z-sections of the nucleus. The nucleus aspect ratio of hiPSC-CMs on soft and stiff unpatterned, as well as glass substrates, remained unchanged, while for patterned soft and stiff substrates, it increased significantly compared to unpatterned and glass. The highest value for the nucleus aspect ratio was observed for cells on micropatterned stiff substrates (Figure 7D).

The normalized area of mitochondria was measured in cells cultured on soft and stiff, patterned and unpatterned substrates, as well as glass (Figure 7E). Our findings indicate a significant increase in mitochondria area when hiPSC-CMs cultured on soft substrate stiffness, consistent with previous studies [51]. On both stiff patterned and unpatterned substrates, mitochondria density decreased dramatically compared to the soft substrates, as illustrated in Figure 7E.

To study the effect of substrate stiffness on hiPSC-CM electrophysiological properties and contractility, cells were grown on both soft and stiff PDMS substrates, with and without patterning. As mentioned earlier, hiPSC-CMs cultured for 7 days on micropatterned substrates represent an immature phenotype compared to those cultured for 22 days. We analyzed Ca^2+^ transients and contractility of hiPSC-CMs at both immature (7 days) and mature (22 days) stages on micropatterned PDMS substrate with different stiffnesses. The results of SarcTrack and OM analysis show that hiPSC-CMs cultured for 7 days on micropatterned stiff substrates exhibit increased contraction and faster rise time compared to those on soft substrates (Figure 8A-D and I-J). Specifically, hiPSC-CMs cultured for 7 days on soft micropatterned substrates showed the highest half-rise time compared to all other conditions (Figure 8A). Contrary to expectations, at day 7 post-replating, no differences observed in the time-to-peak and half-rise time for hiPSC-CMs cultured on soft micropatterned substrates compared to soft unpatterned ones (Figure 8A and B), further indicating the immaturity of hiPSC-CMs at this stage. The fastest half-rise time was observed for cells cultured on stiff micropatterned substrates for 7 days (Figure 8A). The Ca^2+^ transient trace showed a faster decay for hiPSC-CMs cultured on stiff micropatterned substrate compared to soft ones at day 7 post replating (Figure 8D). Additionally, SarcTrack analysis at day 7 revealed a significant increase in contraction velocity for cells cultured on stiff micropatterned substrates compared to soft ones (Figure 8I). Furthermore, contractility traces of hiPSC-CMs showed higher contractility and increased sarcomere shortening in cells cultured on stiff micropatterned substrates at day 7 post-replating (Figure 8J). Consistent results from SarcTrack and OM analysis indicate that after 7 days of replating hiPSC-CMs on soft and stiff micropatterned substrates, the highest contraction and Ca^2+^ transient kinetics were observed for cells on stiff substrate.

**Figure 8.**
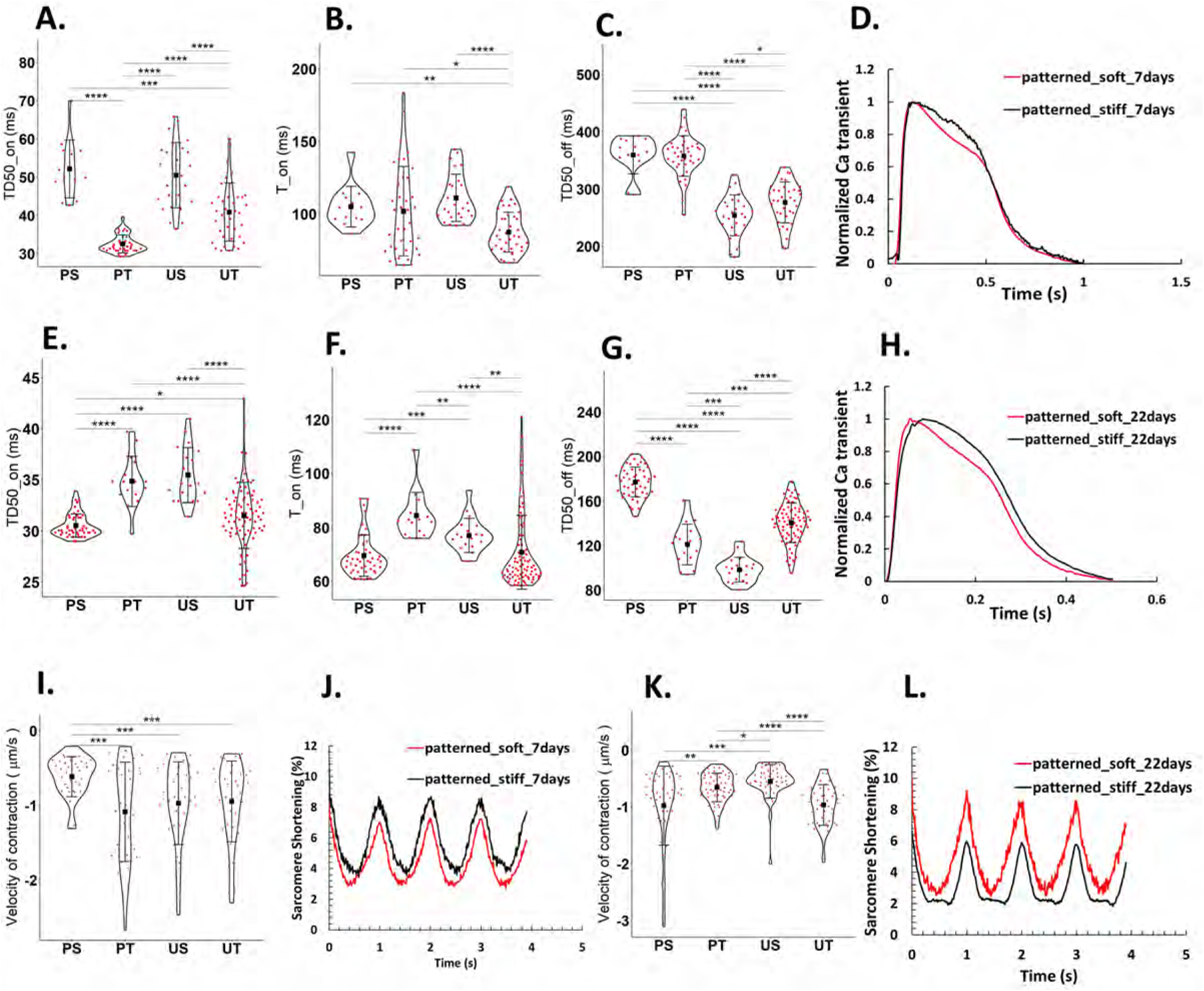
The effect of substrate stiffness on the contractility and electrophysiological properties of hiPSC-CMs at days 7 and 22 post replating. A) Half rise time, B) Time to peak, and C) Half decay time of hiPSC-CMs cultured on stiff and soft PDMS substrate, with and without patterning, at day 7 post replating. D) Normalized Ca transient signal of hiPSC-CMs cultured on soft and stiff micropatterned substrates at day 7. Each Ca transient trace represents the average of at least 120 single traces. E) Half rise time, F) Time to peak, and G) Half decay time of hiPSC-CMs cultured on stiff and soft PDMS substrate, with and without patterning, at day 22 post replating. H) Normalized Ca transient signal of hiPSC-CMs cultured on soft and stiff micropatterned substrates at day 22 post replating. Each Ca transient trace represents the average of at least 120 single traces. I) The velocity of contraction of hiPSC-CMs cultured on stiff and soft PDMS substrate, with and without patterning, at day 7 post replating using SarcTrack. J) The sarcomere shortening trace of hiPSC-CMs cultured on soft and stiff micropatterned substrates at day 7 post replating. Each contractility trace represents the average of at least 40 videos. K) The velocity of contraction of hiPSC-CMs cultured on stiff and soft PDMS substrate, with and without patterning, at day 22 post replating using SarcTrack. L) The sarcomere shortening trace of hiPSC-CMs cultured on soft and stiff micropatterned substrates at day 22 post replating. Each contractility trace represents the average of at least 40 videos. ns p-value > 0.05, * p-value < 0.05, ** p-value < 0.005, *** p-value < 0.0005, **** p-value < 0.00005.

However, results changed significantly after 22 days of culture. The lowest peak time and half-rise time was observed for hiPSC-CMs cultured on soft compared to stiff micropatterned substrates (Figure 8E and F). Moreover, the Ca^2+^ transient decay is dramatically faster in hiPSC-CMs cultured on soft compared to stiff micropatterned substrates (Figure 8H). Additionally, SarcTrack data revealed that increasing substrate stiffness decreased sarcomere shortening and contraction velocity of hiPSC-CMs, as shown in Figure 8K and L, further supporting the OM data at day 22. Furthermore, the effect of patterning on the functional maturation of hiPSC-CMs is evident, as peak time and half-rise time are faster in hiPSC-CMs cultured on soft micropatterned substrates compared to soft unpatterned substrates after 22 days of replating (Figure 8E and F). While the effect of micropatterning on the functional maturation of hiPSC-CMs was not evident at day 7 post-replating, three weeks of culturing the cells on soft micropatterned substrates facilitated maturation.

## 4. DISCUSSION

In this study, we developed a novel micropatterned substrate with varying stiffnesses to explore the structural and functional maturation of hiPSC-CMs over a developmental time course and to mimic both physiological and pathological conditions. The cell culture chamber, equipped with platinum electrodes, facilitated pacing the cells at various frequencies within a controlled environment, ensuring contamination-free conditions. The soft PDMS micropatterned substrate required only 100,000 cells to cultivate a mature monolayer of hiPSC-CMs in 2-4 weeks, with the optimal timing being 3 weeks. Another advantage of this novel substrate is its high reproducibility and low sample-to-sample variability due to the enduring nature of the permanent mold. Additionally, the tunable stiffness of PDMS enables the investigation of various cardiomyopathies and channelopathies responses to fibrotic conditions. Our findings underscore the cost-effectiveness of the cell culture substrate, its independence from clean room facilities, the successful attachment of a 2D monolayer of beating cardiomyocytes after 90 days of culture on the soft substrate, its contamination-free nature, versatility in design with different aspect ratios, and suitability for long-term structural, functional, and electrophysiological analyses.

### 4a. Micropatterned substrate fabrication and surface modification

All soft PDMS substrates were fabricated with consistent specifications, including curing time and spinner speed at each stage, to ensure a uniform thickness range (Figure 1). It is crucial to avoid using a very thin layer of soft PDMS because substrate stiffness changes at lower thickness levels, potentially causing cells to sense the stiff glass at the bottom. Therefore, maintaining consistency in substrate thickness is essential for obtaining reliable and meaningful results when studying the effect of substrate stiffness on hiPSC-CMs maturation and functionality [30][43].

Previous studies have treated the surface of PDMS with PD/Laminin to facilitate the long-term differentiation of hiPSCs into cardiomyocytes, and our results are in line with their findings [26]. PD is derived from the oxidative polymerization of dopamine under alkaline Tris-HCl buffer (pH 8.5) [27]. The adhesive properties of PD enable the formation of a stable layer on PDMS surfaces, acting as a linker to establish covalent bonds with ECM proteins [49]–[51]. The strong covalent bond between the PD-treated PDMS surface and ECM proteins leads to a significant reduction in the contact angle from approximately 90 to 60-40 degrees (Figure 3B). This change in the contact angle facilitates cell adhesion to the substrate, allowing the cells to form a monolayer and exhibit beating behavior. Thus, PD can effectively serve as a linker between PDMS and ECM proteins, enhancing hiPSC-CMs attachment to the substrate without adversely affecting cell functionality. The findings from this study demonstrate that cardiomyocytes on micropatterned PD/Laminin-coated substrates remain attached and functional for more than three months, as evidenced in Supplemental Video 1.

### 4b. Micropatterning enhances the structural and functional maturation of hiPSC-CMs

Fetal to adult development of cardiomyocytes leads to a transition to rod-shaped morphology characterized by organized and aligned sarcomeres, increased percentage of binucleated hiPSC-CMs, enhanced nucleus aspect ratio, increased mitochondria density favoring fatty acid oxidation over glycolysis, increased contractility, and improved calcium transient kinetics facilitated by gap junction formation [52][52]. The results of our study revealed an enhancement of many of these structural and functional maturation markers in hiPSC-CMs cultured on a soft micropatterned substrate.

Our results demonstrated that culturing hiPSC-CMs on a micropatterned PDMS substrate for 22 days significantly enhances maturation, as evidenced by: 1) Improved sarcomere organization and orientation (Figure 4A-C), 2) An increase in the nucleus aspect ratio (Figure 4D), 3) Enhanced mitochondria density (Figure 7E) 3) Increased sarcomere length (Figure 5), 4) Accelerated contraction and relaxation kinetics (Figure 6F and G), 5) Higher sarcomere shortening (Figure 6E), 6) Faster calcium release and reuptake (Figure 6A-D), 7) Enhanced calcium transient amplitude (Figure 6C), and 8) restitution in response to high pacing frequencies (Figure 6H-L).

While the highest sarcomere length and contractility observed after three weeks, the lowest sarcomere length detected after 3 months of culturing hiPSC-CMs on soft patterned and unpatterned substrates (Figure 5). This shift can be explained through the dedifferentiation, proliferation, and likely redifferentiation of adult cardiomyocytes, as suggested in previous research [53]. Cardiomyocyte dedifferentiation and redifferentiation can be identified through gene expression and functional analysis, including sarcomeric organization and structure [53]. Previous studies have demonstrated that coculturing adult cardiomyocytes isolated from mice with neonatal rat ventricular myocytes (NRVMs) led to myocardial dedifferentiation, proliferation, and redifferentiation (DPR) into functional adult cardiomyocytes through gap junction formation between new cardiomyocytes and NRVMs [54]. Based on our results shown in Figure 7A and B, cardiomyocytes exhibited a more adult-like phenotype after 3 weeks of culturing on a micropatterned substrate with 20 µm (C-C) and depth of D2. However, at day 30, the sarcomere length slightly decreased compared to day 20, possibly indicating the initiation of the dedifferentiation process (Figure 5).

As depicted in Supplementary Video 3, a 2D monolayer of cardiomyocytes at day 20 transformed into a 3D thick multilayer of cardiomyocytes with thickness changes approximately from 10 to 25 µm, through days 20 to 90, respectively. This transformation may represent the proliferation of dedifferentiated cardiomyocytes due to hypoxia or stress in the environmental condition. The bottom layer of hiPSC-CMs on the micropatterned soft substrate was aligned at day 90, while the sarcomeres at the top layers were disorganized and resembled the immature cardiomyocyte phenotype. Therefore, the disorganized sarcomeres at the top layer of aligned cardiomyocytes after 90 days on the micropatterned substrate could represent two important processes: 1) the proliferation of dedifferentiated cardiomyocytes, or 2) redifferentiation of hiPSC-CMs due to gap junction formation between redifferentiated hiPSC-CMs and spontaneously beating hiPSC-CMs. However, further studies are required to confirm the DPR process of hiPSC-CMs on soft PDMS substrates.

### 4c. Stiffness influence on cellular orientation, nuclei aspect ratio, mitochondrial density, contractility, and Ca^2+^ transients

As substrates become stiffer, their ability to align the cells enhances significantly, due to less movement and more stability of substrate during each beat compared to softer substrates. This slightly facilitates the process of cardiomyocyte and nucleus alignment and elongation in the direction of the patterns. Hence, no significant differences in nucleus aspect ratio, sarcomere organization, and OOP were detected for hiPSC-CMs cultured on soft compared to stiff micropatterned substrates. However, mitochondria density changed significantly when culturing on substrate with different stiffnesses which is consistent with previous studies [55]. Mitochondria occupy approximately 30% of adult native cardiomyocyte volume. In immature cardiomyocytes, mitochondria exhibit a circular morphology and are more sparce [51]. As cells mature, mitochondria area intensity significantly enhances, leading to an elongated configuration aligned with the myofibril structure. When stiffness is elevated towards pathological levels, mitochondria revert to a fetal phenotype due to cardiac remodelling leading to lower mitochondrial content [55]–[57].The percentage of mitochondria within a 2D monolayer of hiPSC-CMs cultured on a soft micropatterned substrate was approximately 18%, as illustrated in Figure 7E. In contrast, the lowest reported value was observed in hiPSC-CMs on glass substrates, measuring about 10%. Consequently, the use of micropatterned soft PDMS substrates led to an approximately two-fold increase in mitochondrial content compared to cells cultured on glass dishes.

Substrate stiffness significantly affect the contractility and Ca^2+^ transient during developmental time course. Consistent results from SarcTrack and OM analysis indicate that after 7 days of replating hiPSC-CMs on soft and stiff micropatterned substrates, the highest contraction and Ca^2+^ transient kinetics were observed for cells on stiff substrate. The observed increased contractility and kinetics of Ca^2+^ transients of hiPSC-CMs on stiff substrate at day 7 post-replating compared to the soft substrate could be attributed to factors such as the immaturity of hiPSC-CMs or remodeling observed during fibrosis. However, after 22 days, hiPSC-CMs exhibited a mature phenotype, aiding in representing the phenotype of the cells when cultured on both soft and stiff substrates. Notably, hiPSC-CMs cultured on stiff micropatterned substrates for 22 days exhibited decreased sarcomere shortening, slowed rise time and rate of Ca^2+^ removal, and altered Ca^2+^ homeostasis as expected in fibrosis. The observed increase in sarcomere shortening, contractility, and calcium transient kinetics at day 22 post-replating of hiPSC-CMs on a soft micropatterned substrate aligns with the heightened mitochondria density, indicative of elevated energy demand.

### 4d. Limitations and future directions

This study primarily aimed to develop a soft micropatterned substrate to promote the maturation and enable long-term structural and functional analysis of 2D monolayer hiPSC-CMs. However, the utilization of this substrate for aligning and maturing cardiomyocytes in 3D structures has not been explored. Additionally, the impact of the micropatterned soft substrate on protein and gene expression profiles was not investigated. In future studies, we intend to address these limitations by employing techniques such as printing cardiomyocytes on the soft micropatterned substrate to examine the effect of patterning on the alignment of hiPSC-CMs embedded in a hydrogel. Furthermore, mass spectrometry will be utilized to assess the protein expression of hiPSC-CMs on micropatterned PDMS substrates.

## 5. CONCLUSIONS

In conclusion, this study utilized PDMS to establish a physiologically relevant environment for the maturation of hiPSC-CMs. The introduction of a soft micropatterned substrate enabled hiPSC-CMs to form a 2D monolayer. Notably, for the first time, we demonstrated that surface modification with PD/Laminin did not adversely affect cell functionality and facilitated robust cell attachment on soft substrates for an extended period exceeding three months. To explore the optimal pattern dimensions for achieving the highest level of maturation, various pattern dimensions were examined. It was observed that the 8 to 10 µm depth patterns were ineffective in aligning hiPSC-CMs. In contrast, a 2.5-5 µm deep and 20 µm (C-C) pattern with 2-5 kPa stiffness, proved to be the most effective. This pattern dimension demonstrated superior alignment and elongation of hiPSC-CMs in the direction of the pattern, a more uniform OOP distribution, an increase in the nucleus aspect ratio, enhanced mitochondria density, enhanced contractility, and improved kinetics of Ca^2+^ transient. For the first time, we conducted a comprehensive assessment of cell contractility over an extended culture duration of approximately three months on soft micropatterned substrates. This investigation aimed to identify the optimal time frame during which hiPSC-CMs reach the highest level of maturation. As our samples were equipped with electrodes in a closed chamber system, we monitored the same sample at intervals of 3 to 90 days after replating cells onto the substrates. The maximum resting sarcomere length, exceeding 2 µm was observed after three weeks of culturing the cells on the soft micropatterned substrates with a 20 µm (C-C) and a depth of 2.5-5 µm, indicating hiPSC-CMs maturation. Additionally, the kinetics of Ca^2+^ transient revealed faster rise time and half decay time when cells were cultured on soft micropatterned substrate for 22 days. Furthermore, the investigation delved into the influence of substrate stiffness on hiPSC-CM contractility and Ca^2+^ transient. Results revealed a significant reduction in contractility and a slowed rise time when cells were cultured on stiff micropatterned substrates with a stiffness of 50 kPa for 22 days, which recapitulates a fibrotic condition. These findings contribute valuable insights into optimizing the microenvironment for hiPSC-CM maturation, offering implications for the advancement of cardiac tissue engineering and disease modelling. The developed methodologies and identified parameters represent crucial steps forward in ongoing efforts to enhance the functionality and maturity of hiPSC-CMs, holding significant promise for applications in regenerative medicine and drug discovery.

## Abbreviations

PDMS: Polydimethylsiloxane
hiPSCs: Human induced pluripotent stem cells
hiPSC-CMs: Human induced pluripotent stem cell-derived cardiomyocytes
PD: Polydopamine
C-C: Center to center dimension of the grooves
D1: Depth of grooves of ranging from 8 to10 μm
D2: Depth of grooves of ranging from 2 to 5 μm
PVA: Polyvinyl alcohol
PS: Patterned soft PDMS substrate
PT: Patterned stiff PDMS substrate
US: Unpatterned soft PDMS substrate
UT: Unpatterned stiff PDMS substrate

## Supplementary File

### OOP Image Analysis Workflow

To determine the Orientational Order Parameter (OOP), two types of images were utilized: SP8 Laser Scanning Confocal Microscope images at 40X and 63X magnifications and spinning disk images at 60X with water immersion. Confocal images were cropped to a resolution of 1024x1024 pixels while Nikon spinning disk images were maintained at their original resolution of 1626x1160 pixels. The image processing workflow for measuring the OOP is shown in Figure S1. To elucidate the workflow, two distinct programs were utilized. The first program, GTFiber, is a MATLAB package designed to quantify fibre alignment. The second program, CellProfiler, was utilized for filtering out objects that are not suitable for our application. The parameters of the GTFiber package for images acquired from a confocal spinning disk are: Gaussian smoothing, 0.5 µm orientation smoothing, 1 *μm*; diffusion time, 5 s; top hat size, 1 µm aptive thresholding; noise removal, 0.5 *μm*^2^.

After thresholding and image cleaning, a custom CellProfiler pipeline is employed to filter objects based on MaxAxisLength (10-120 pixels) and Orientation parameters. This step aims to eliminate objects that are perpendicular to the Z-Lines and exhibit long fibre sizes. Finally, the output of the CellProfiler workflow is used to measure the OOP with the GTFiber package. In this study, the grid step of 20 µm and frame step of 10 are used to determine the *S*_*full*_ as the OOP of the Z-Lines.

Figure S2 illustrates the image processing workflow for the captured images from the confocal spinning disk or SP8 laser scanning confocal microscope depicting α-actinin (green) and nuclei (blue) of fluorescent-tagged hiPSC-CMs after 7 days of replating. The initial step involves the utilization of the GTFiber package in MATLAB, which incorporates coherence-enhancing anisotropic diffusion filtering and top-hat filtering. After image thresholding and cleaning, off-targeted fibres persist in the GTFiber skeleton image output.

Figure S2B shows the skeleton images of fibers detected only by the GTFiber output. In Figure S2C, the orientational map reveals off-target fibers that are perpendicular to the z-lines or have orientations misaligned with the orientation of z-lines. To address this issue, a custom CellProfiler workflow is implemented to filter out off-target fibers detected by the GTFiber package. This workflow is directly applied to 2D hiPSC-CMs monolayers cultured on micropatterned substrates, employing two object filters: MajorLengthAxis and orientation (Figure S2E, S2F).

The results in Figure S2H demonstrate the successful removal of off-target fibers, leaving only those representing the Z-lines of hiPSC-CMs. For hiPSC-CMs cultured on unpatterned and glass substrates, the same workflow is implemented, but only the MajorLengthAxis parameter in CellProfiler is chosen for filtering.

### Nucleus Image Analysis Workflow

The nucleus was identified using a custom CellProfiler pipeline. The blue, fluorescent channel (DAPI) is used for nucleus analysis. Image preprocessing methods, including Gaussian and Median filters, as well as the Sobel Edge-finding method, were employed to enhance the detection of the nucleus. In the pipeline, adaptive Minimum Cross-Entropy was employed to segment DAPI images, facilitating the identification of nucleus.

### Mitochondria Image Analysis Workflow

In our mitochondria analysis, encompassing Z-line detection and determination of the OOP, we employed the GTFiber and CellProfiler programs, as shown in Figure S4 The GTFiber was configured with specific parameters optimized for images obtained from a confocal spinning disk, Gaussian smoothing, 0.5 *μm* ; orientation smoothing,1 µm ffusion time,5 s; top hat size, 1 *μm* ; adaptive thresholding; noise removal,0.5 *μm*^2^. The resulting images from GTFiber were processed through a custom CellProfiler pipeline to identify mitochondria as the primary object and measure their pixel areas.

As our images are derived from a 2D monolayer of hiPSC-CMs, covering the entire image area with cells, we normalized the number of pixels associated with the mitochondria area by the total number of pixels in each image. For Nikon spinning disk images, a consistent resolution of 1626x1160 pixels was maintained. The parameter derived from this normalization process facilitates the comparison of different conditions in hiPSC-CMs cell culturing.

### Ca^2+^ transient temporal parameters

All the temporal parameters of the Ca^2+^ transient are listed below and depicted in Figure S5.

T0 : Time obtained as the intersection of the trace (before peak) and baseline.

Tend: Time obtained as the intersection of the trace (after peak) and baseline.

Tpeak: peak time of the fluorescence trace.

Baseline: Avarage value of the trace that corresponds to a temporal window of 20% of the pacing period. It is also considered as F0.

Fmax/F0: maximum flourecent value (trace value) divided by baseline value.

T_on: time obtained from T0 until the peak value of a trace.

T_off: time from peak to the Tend.

TD50_on: obtained from T0 and time to 50% contraction.

TD50_off: Time from peak to 50% relaxation.

---

**Figure S1.**
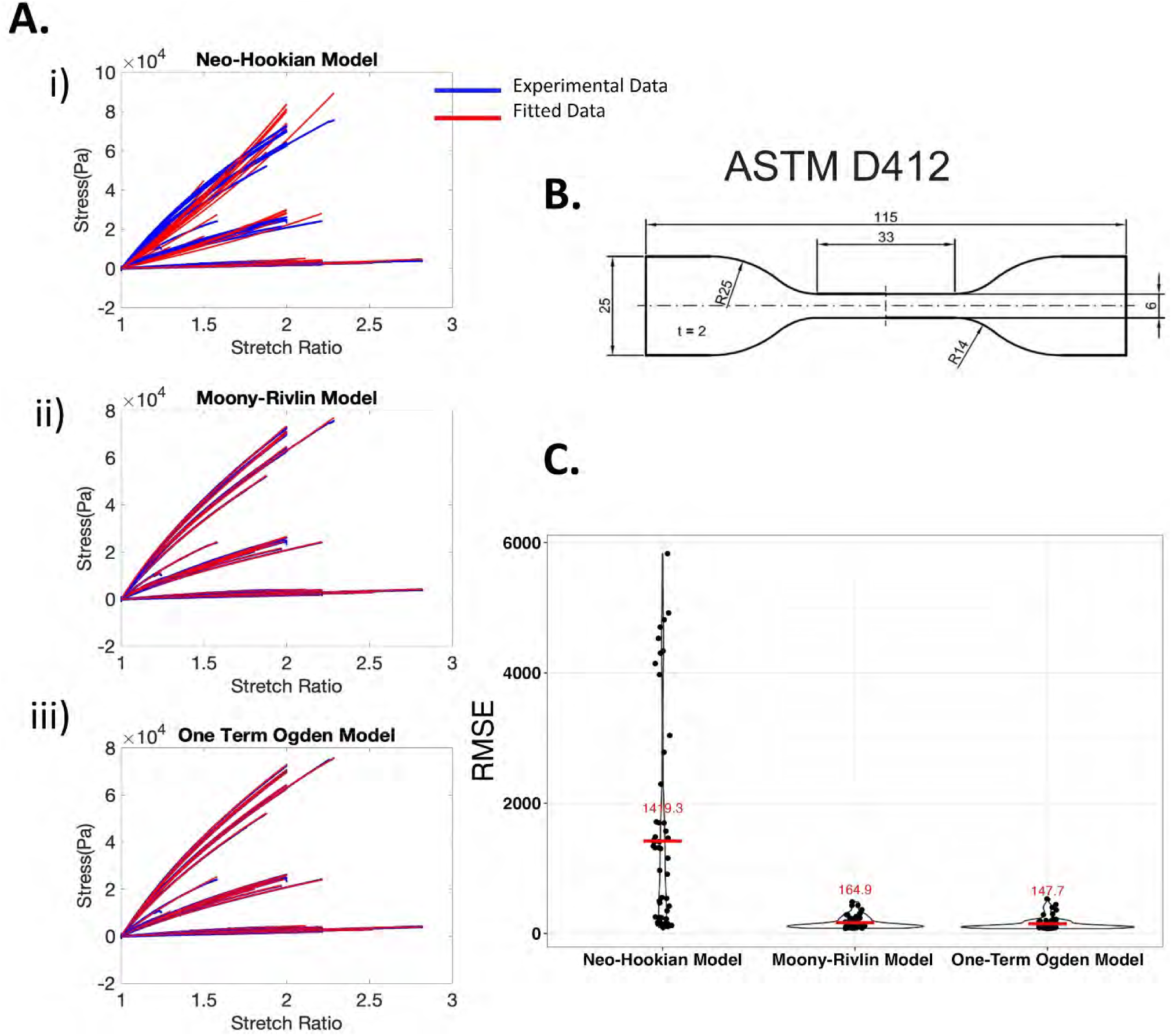
The uniaxial test results and using three models to fit the experimental data: (A-i) Neo-Hookian Model (A-ii) Moony-Rivlin Model (A-iii) One-Term Ogden Model (B)Uniaxial test sample (ASTM D412) (C) RMSE criteria.

**Figure S2.**
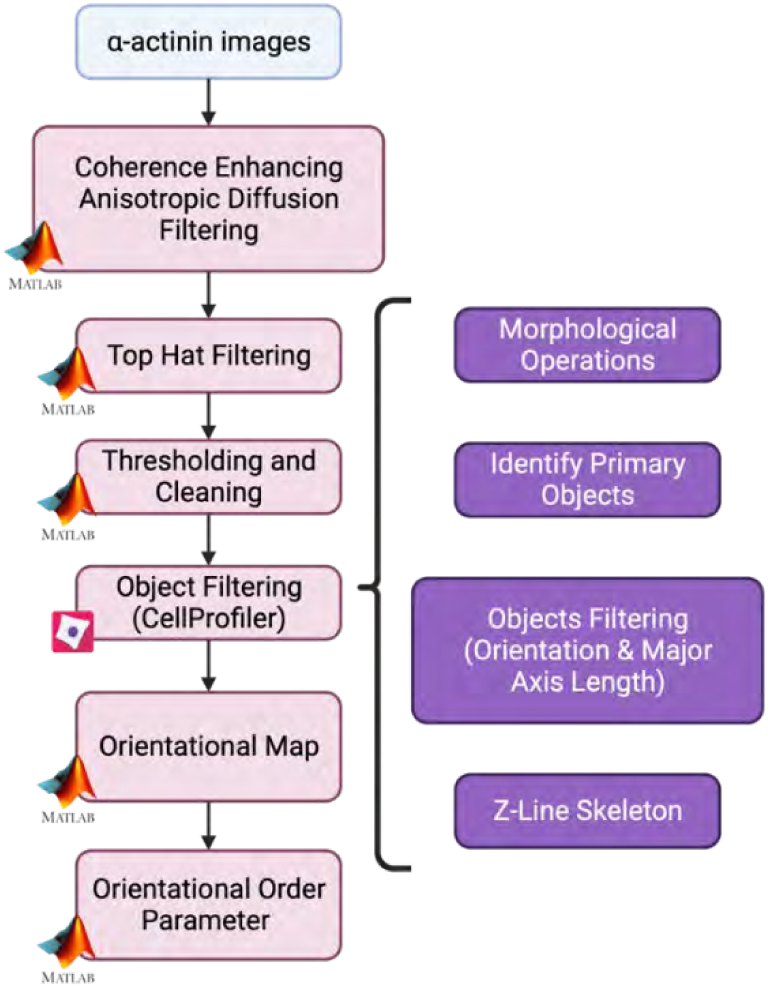
Outline of the image processing workflow used to determine the OOP of Z-Lines.

**Figure S3.**
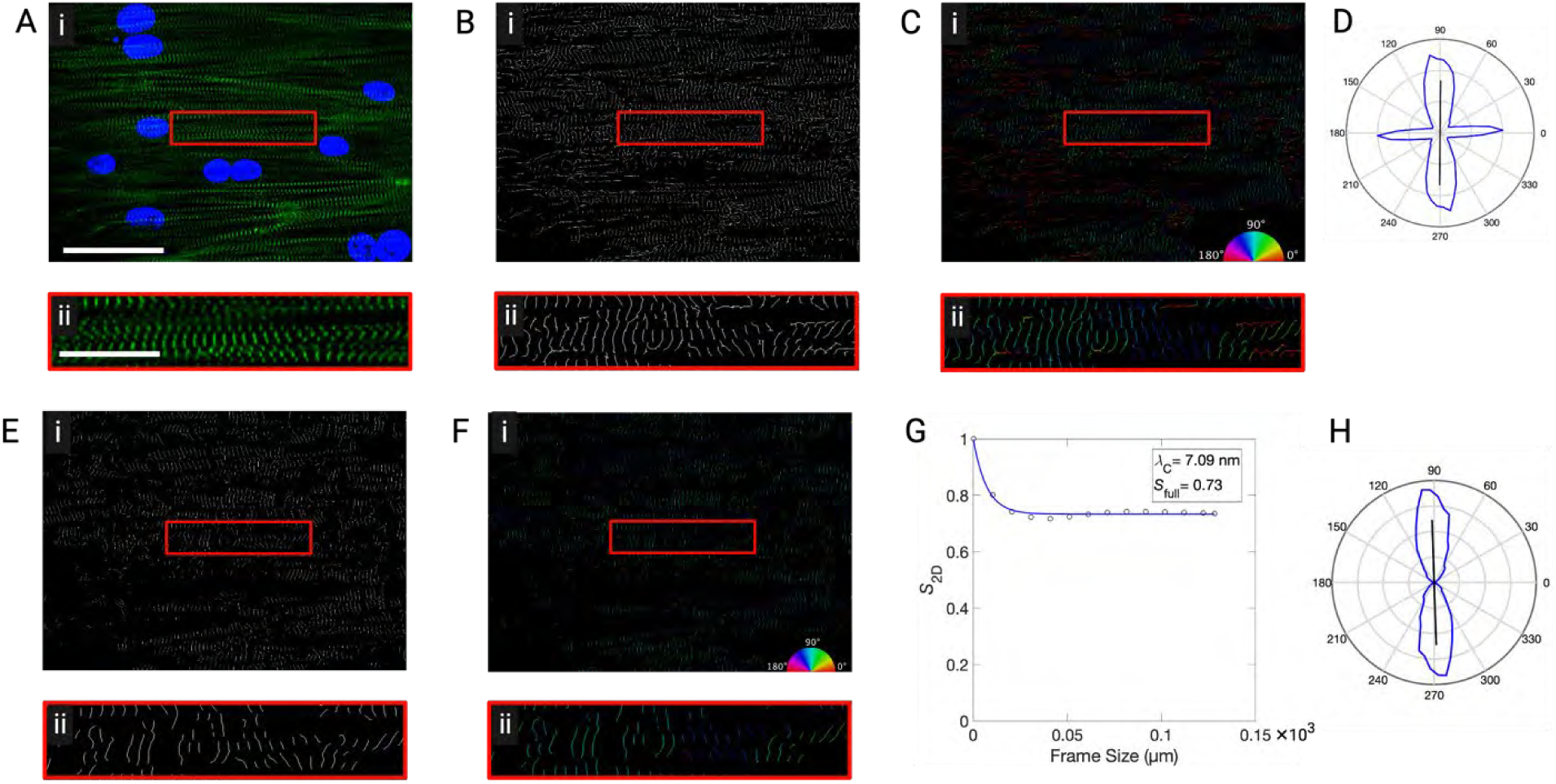
Example images representing the procedure used to determine OOP of Z-Lines in hiPSC-CMs. (A)The confocal spinning disk images of a-actinin (green), and nuclei (blue) of fluorescent-tagged hiPSC-CMs (After 7 days replating). (B) The Skeleton image shows the skeletonization of a-actinin by using the GTFiber, a MATLAB program. (C) colour Orientational Map using the GTFiber. Each pixel’s orientation corresponds to an orientation on the attached colour wheel. (D) Orientational Distribution extracted from Orientational Map. (E) Skeleton image output extracted from CellProfiler pipeline. (F) Orientational Map for skeleton image of CellProfiler pipeline. (G) Decay of the OOP, *S*_*2D*_, as a function of frame size. Fitted model parameters are indicated at the upper right. The *S*_*full*_, the asymptotic value of *S*_*2D*_ is considered as OOP in this study. (H) Orientational Distribution extracted from the Orientational Map after using the CellProfiler pipeline to filter objects. Scale bars: (A-C i) 50 *μm*; (A-C ii) 20 *μm;*; (E-F i) 50 *μm*; (E-F ii) 20 *μm*.

**Figure S4.**
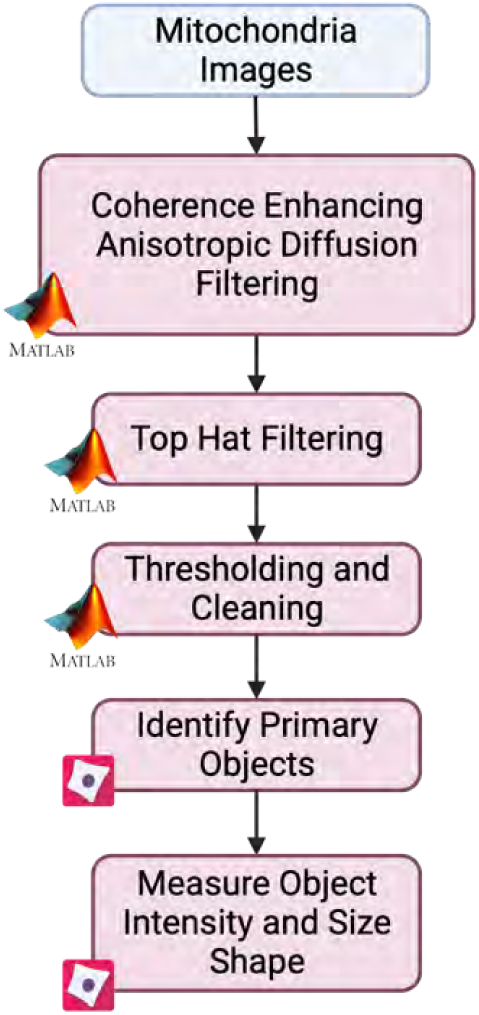
The workflow for mitochondria analysis.

**Figure S5.**
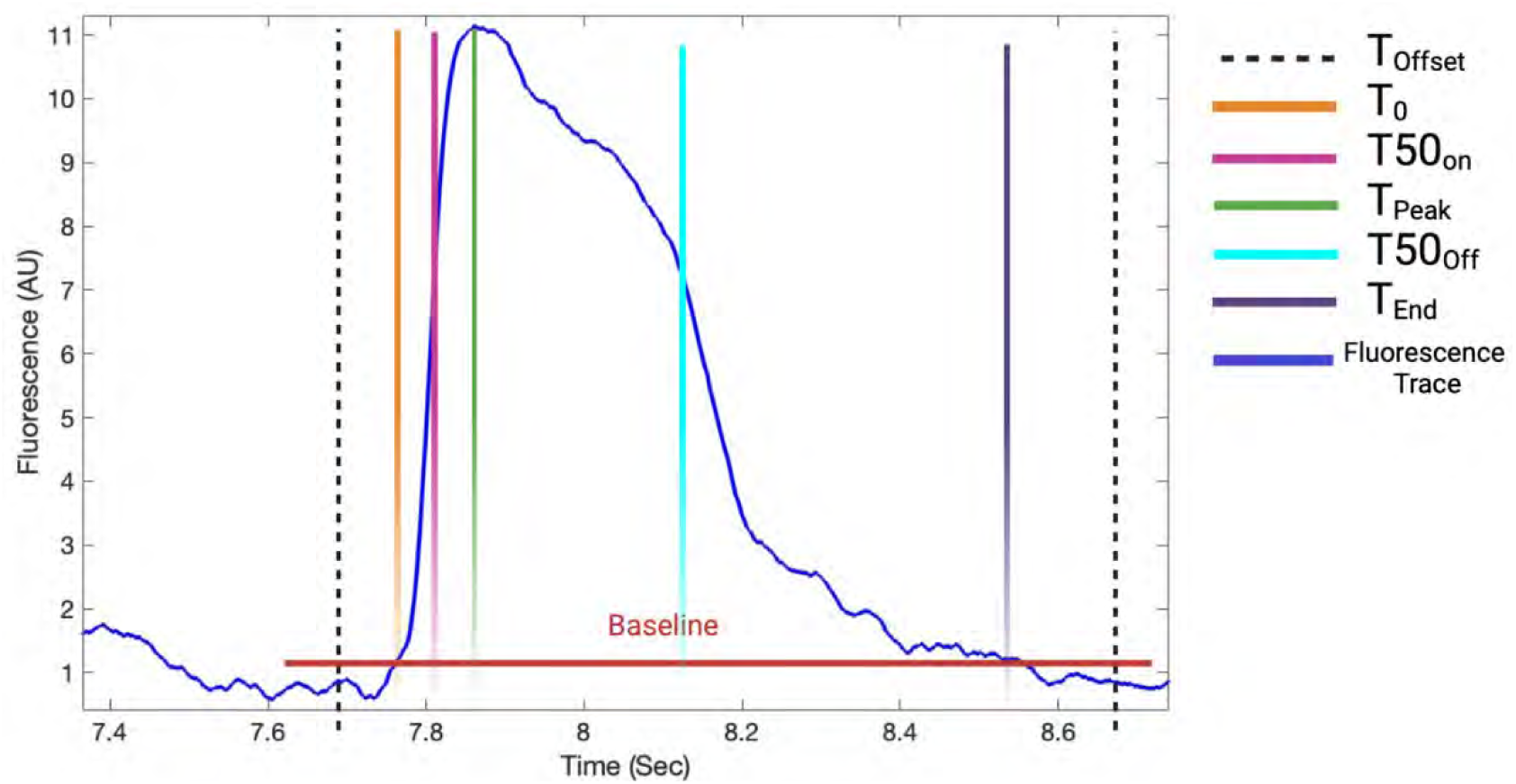
The temporal parameters of Ca^2+^ transient.

**Figure S6.**
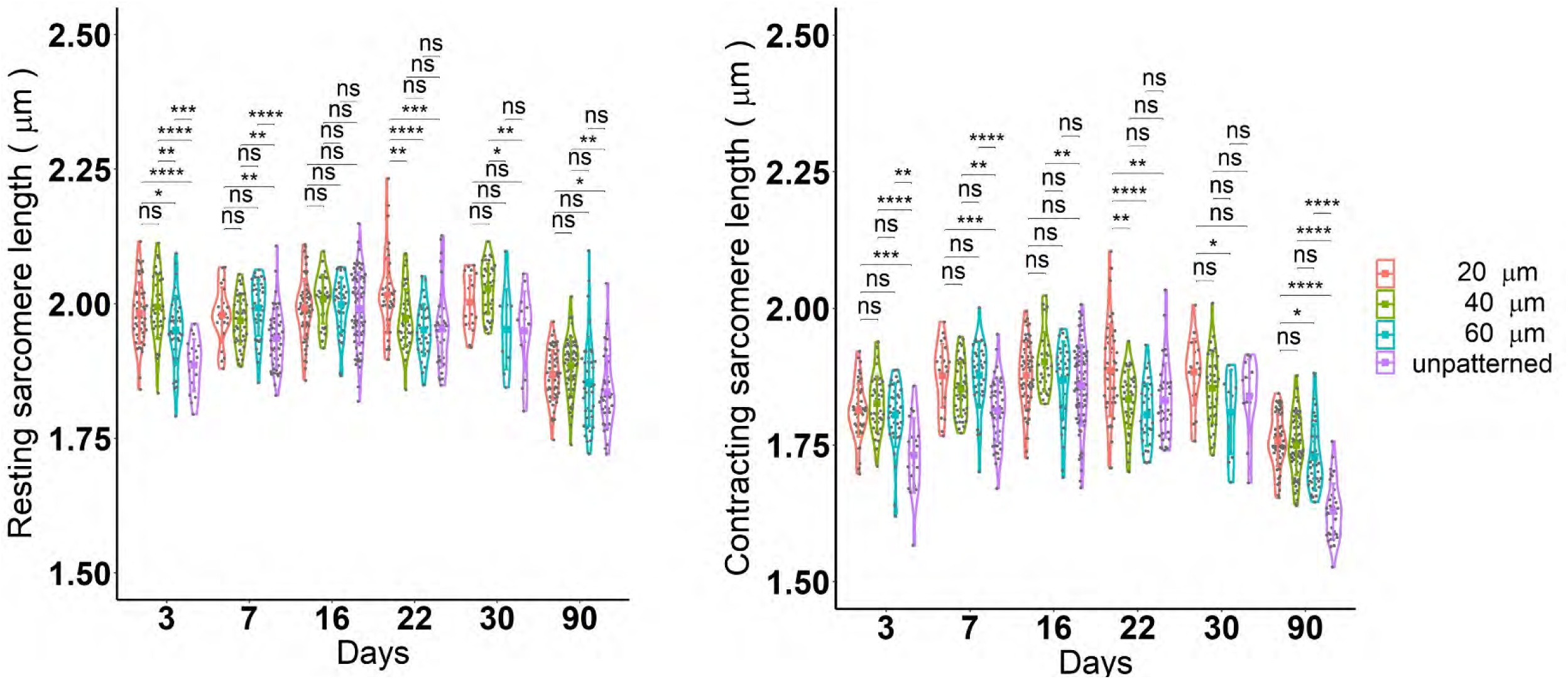
The resting and contracting sarcomere length of hiPSC-CMs cultured on unpatterned (purple) and micropatterned PDMS substrates with different pattern dimensions of C-C of 20 µm (red), 40 µm (green), and 60 µm (blue) with a depth of 5 μm at intervals of 3, 7, 14, 22, 30, and 90 days post-replating. A pairwise t-test with Fdr adjustment for p-value was used to determine the significant level between different pattern sizes at each day. ns p-value > 0.05, * p-value < 0.05, ** p-value < 0.005, *** p-value < 0.0005, **** p-value < 0.00005.

## Notes

### Competing Interest Statement

The authors have declared no competing interest.

